# Whole brain dimensional approach identifies networks of stress susceptibility in male and female mice

**DOI:** 10.1101/2025.05.10.653278

**Authors:** Lizette Herrera-Portillo, Daniel Gallino, Yohan Yee, Jessie Muir, Gabriel A. Devenyi, Rosemary C. Bagot, M. Mallar Chakravarty

## Abstract

Stress is a major risk factor for depression and anxiety, two highly comorbid disorders with sex differences in symptom presentation and prevalence. Accumulating evidence on the chronic variable stress (CVS) mouse model has revealed that stress susceptibility is sex-specific, as females require less stress exposure (6 days) compared to males (21-28 days) to develop comparable adaptations in depressive- and anxiety-like behaviors. However, the neuroanatomical changes associated with stress susceptibility remain unclear in each sex. Given this differential susceptibility, elucidating these mechanisms requires a phenotype-matched approach in which stress duration is titrated in a sex-specific manner to induce comparable behavioral adaptations. Here, we used a multi-variate brain-behavior and connectomic analysis to examine signatures of stress susceptibility in female and male mice. Using structural magnetic resonance imaging (MRI), we investigate whole-brain neuroanatomical changes associated with behavioral susceptibility within each sex. We then examined the structural connectome underlying neuroanatomical changes associated with behavioral susceptibility within each sex, and putative molecular basis for behaviorally-relevant changes using spatial gene expression analyses. CVS induced significant neuroanatomical changes in regions commonly implicated in anxiety- and depressive-like phenotypes in both sexes (e.g. nucleus accumbens and hippocampus); however qualitative differences between sexes in the direction of change were observed. In addition to these changes, in females stress-induced neuroanatomical changes were associated with both depressive- and anxiety-like behavior, whereas in males we observed two orthogonal dimensions of neuroanatomical changes associated with anxiety-like behavior or social preference. The connectomic topology underlying these brain-behavior dimensions revealed the cortical hub regions for females and subcortical hub regions for males. In females, neuroanatomical changes were associated with genes enriched for protein localization to the cell surface, suggesting alterations in cholinergic signaling By implementing a phenotype-matched CVS protocol, our findings indicate that the neuroanatomical signature of stress susceptibility encompasses similar brain regions across both sexes, however the direction of change, association to behavior and connectomic properties may differ in each sex.

## 1. Introduction

Depressive and anxiety disorders are among the leading causes of disease burden worldwide (GBD 2019 Mental Disorders Collaborators, 2022). These highly comorbid disorders manifest differently across individuals, especially between men and women. In depression, sex or gender differences are observed in the prevalence (Kessler et al., 1993), symptom profiles (Kim et al., 2015) and comorbidities (Zhou et al., 2017). These differences in prevalence cannot be fully explained by the sensitivity or rate of exposure to risk factors like stressful life events (Kendler et al., 2001). Many lines of inquiry suggest that much of the variation in depression- and anxiety-related neural phenotypes observed through magnetic resonance imaging (MRI) (Mohammadi et al., 2023) may result from sex-specific responses to stressful experiences (Goldfarb et al., 2019).

Modeling the impact of chronic stress exposure using rodent models has provided a window into the mechanisms of stress vulnerability in subsets of animals, often classified as “susceptible” or “resilient” based on depressive-like behaviors (Akil & Nestler, 2023). Yet, these categorical labels typically rely on a single behavioral outcome, and animals classified as resilient by one measure can display behavioral adaptations similar to those of susceptible animals across other domains, suggesting that stress adaptations exist along a continuum (Akil & Nestler, 2023). Characterizing this spectrum of stress-induced adaptations using dimensional approaches may reveal phenotypic variability that is overlooked by binary classifications.

The chronic variable stress (CVS) rodent model for depression, in which mice are repeatedly exposed to inescapable stressors, has revealed sex differences in both the time course and mechanisms of susceptibility. Studies exposing both sexes to 6 days of stress show that females, but not males, exhibit behavioral alterations, which are associated with epigenetic mechanisms, such as DNA methyltransferase 3a (Dnmt3a) regulation in the nucleus accumbens (Hodes et al., 2015), and increased excitability of the ventral hippocampus to nucleus accumbens pathway (Williams et al., 2020). Conversely, extending the duration of exposure to 21-28 yields comparable behavioral alterations in males (Labonté et al., 2017; Liu et al., 2018; Muir et al., 2020). While these studies have been instrumental in identifying mechanisms that differentiate males and females under identical stress conditions, they are limited in their ability to examine mechanisms of stress susceptibility within each sex, as similar stress durations induce distinct phenotypes across sexes. Disentangling mechanisms of susceptibility within each sex requires matching the phenotype rather than the duration of exposure, which is achieved by titrating stress exposure in a phenotype-matched manner.

Non-invasive neuroimaging techniques, like small animal magnetic resonance imaging (MRI), provide a valuable opportunity to examine longitudinal changes in neuroanatomy at the whole-brain level in the context of stress susceptibility. Contextualizing these phenotypes within structural connectivity of the mouse brain (Knox et al., 2019; Oh et al., 2014) allows us to situate them within distributed brain circuits. Within this framework, graph theory metrics provide a powerful framework to characterize psychiatrically-relevant properties of brain networks organization (Fornito et al., 2015). Efficient transmission of information between subsystems, or communities, is mediated by densely connected hub regions often referred to as rich-club nodes (Bullmore & Sporns, 2012; Gordon et al., 2018). These hubs sit at the interface between local and global communication dynamics, and represent potential target regions for therapeutic interventions to modulate broader brain networks (Gordon et al., 2018; Horn & Fox, 2020).

Building on previous research in the CVS model from our group (Muir et al., 2020), we conducted a sex-specific investigation of stress susceptibility by examining changes in local brain volume, measured with structural MRI (Figure 1A-B), and in depressive- and anxiety-like behaviors in female and male mice exposed to 6 and 28 days of CVS, respectively. Using multivariate techniques, we derived latent dimensions of maximally covarying patterns of stress-induced changes in neuroanatomy and behavior within each sex (Guma et al., 2021; McIntosh & Lobaugh, 2004) (Figure 1C). We then explore the topology of the structural connectome underlying these latent dimensions using graph theory metrics and the voxel-wise structural brain connectivity data from the Allen Institute for Brain Science (Knox et al., 2019; Oh et al., 2014) to identify putative regional targets that may ameliorate this phenotype (Figure 1D). Finally we used whole-brain spatial gene expression data from the Allen Institute for Brain Science (Lein et al., 2006), to probe potential molecular mechanisms for region-specificity (Figure 1E).

**Figure 1.**
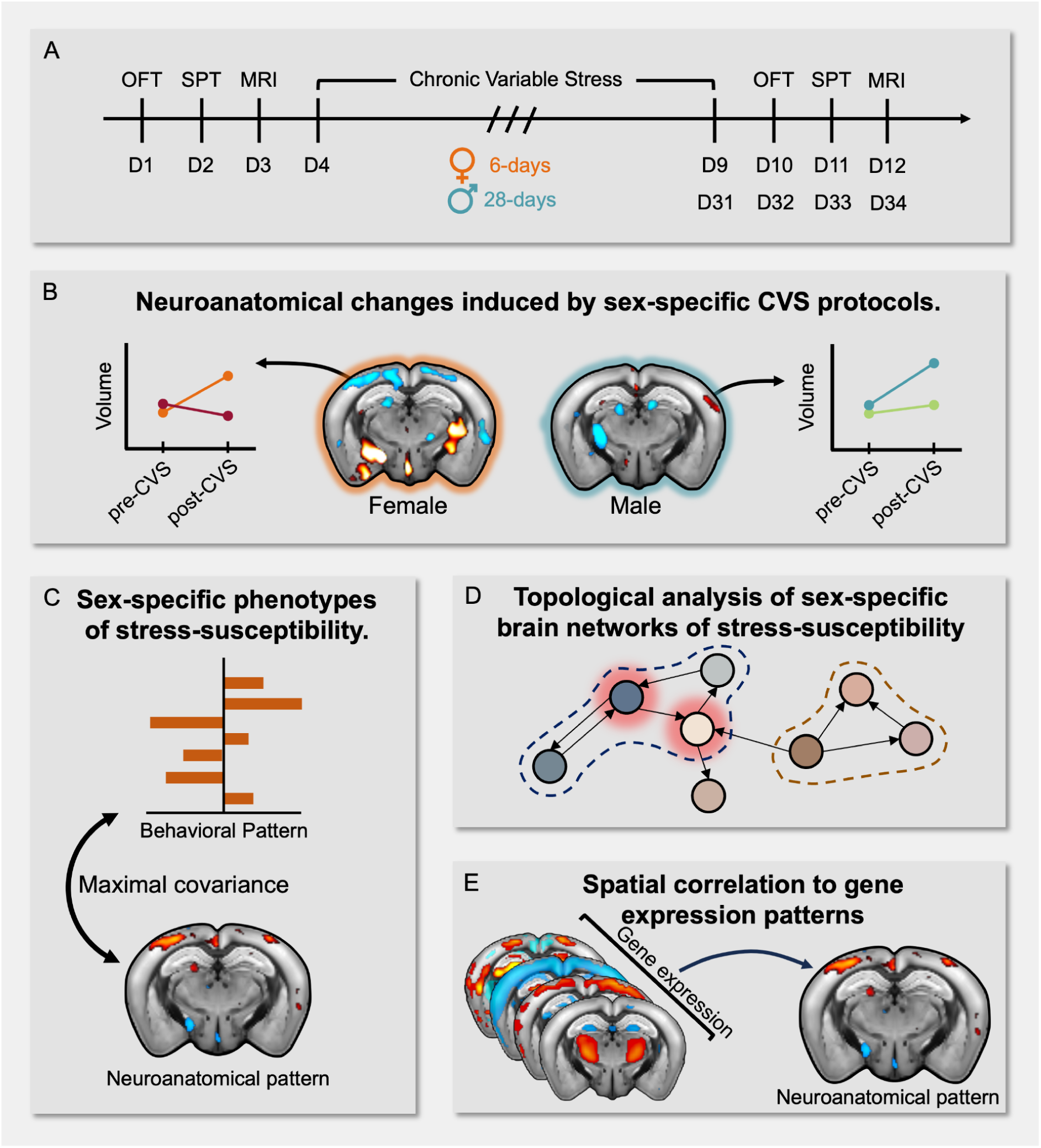
Experimental timeline and methodology. A) Experimental timeline: Behavioral tests of anxiety- (open field test) and depressive-like behaviors (social preference test) and MRI scans were acquired at baseline and 24 hours after the last stressor. To induce behavioral susceptibility, females and males were exposed to 6 and 28 days of CVS, respectively. B) We performed univariate longitudinal analyses on behavioral variables and Jacobian determinants (voxel-wise volume differences) to investigate changes induced by phenotype-matched protocols of CVS. C) To investigate underlying mechanisms of phenotypes of stress-susceptibility in each sex, we examined the correlation between neuroanatomy and behavior using partial least squares correlation (PLSC). This analysis was performed on: within-subject measures of change across time, and 2) cross-sectionally at post-CVS timepoint. D) To examine the topology of the connectome underlying structurally connected regions from our brain-behavior phenotypes we used graph theory analyses. E) To examine potential biological mechanisms underlying stress-susceptibility phenotypes in each sex, we investigated the spatial correlation to gene expression patterns, followed by gene enrichment analyses.

In both sexes, we identified that neuroanatomical alterations were not confined to corticolimbic regions but were also observed in somatosensory and motor areas. In females, we identified a shared phenotype of stress susceptibility in which neuroanatomical changes covary with both depressive- and anxiety-like behaviors. Conversely, in males, we discovered two orthogonal dimensions of neuroanatomical changes associated with depressive- or anxiety-like behaviors. The topology of the structural connectome underlying these latent dimensions revealed rich club nodes identified in just one sex that mediate changes in both local and global communication. Our results underscore the importance of tailoring experimental designs to match the phenotype when investigating sex-specific phenomena, as similar phenotypes may arise from distinct manipulations.

## 2. Results

To examine whole-brain neuroanatomical signatures of stress susceptibility in a sex-specifc manner we exposed females and male mice to 6 days and 28 days of CVS, respectively, based on the work of our group and others (Johnson et al., 2021; Muir et al., 2020). Given these experimental design choices, all analyses were conducted in each sex and the term “sex-specific” is used to denote effects that are qualitatively observed in one sex but not the other. We first report our findings related to stress susceptibility in females, followed by those from our analyses in males.

### 2.1 Behavioral and neuroanatomical changes of female susceptibility

Building on our previous work on sex-specific stress exposure (Muir et al., 2020), female mice were exposed to 6 days of CVS and underwent structural MRI scanning (T1-weighted (T1w) structural images; 100um^3^ voxels; TR: 20ms; TE: 4.5ms; 7T Bruker Biospec 70/30 equipped with a Bruker 1H Quadrature Transmit/receive MRI CryoProbe) before and after CVS to probe neuroanatomical change. Using group-by-timepoint interactions in linear mixed effects models, we examined longitudinal changes as estimated by the group slopes induced by CVS in anxiety- and depressive-like behaviors, and whole-brain neuroanatomy (Figure 1A).

Six days of CVS increased anxiety-like behavior in females, as measured by reduced time spent in the center of the open field arena (Figure 2A) and decreased social preference (fewer social interactions with a novel mouse) (Figure 2C) following stress exposure. By leveraging the non-invasiveness and spatial coverage of MRI, we then examined voxel-wise neuroanatomical changes (estimated using the Jacobian determinant after nonlinear registrations (Chung et al., 2001)) from T1w structural MRI scans using deformation-based morphometry (DBM) (Germann et al., 2025) as we have done in recent publications (Cupo et al., 2025; Guma et al., 2021; Tullo et al., 2025). Linear mixed effects models identified multiple voxels with significant differences in relative volume change across time between stress and control female mice that survived 5% False Discovery Rate (FDR) (Benjamini & Hochberg, 1995) (Figure 2E). The most prominent significant differences included volume increases in the nucleus accumbens, ventral hippocampus, medial amygdala, lateral hypothalamus and lateral septal complex, alongside volume decreases in somatomotor areas, retrosplenial area, nodulus, agranular insula and anterior cingulate area (Supplementary Figure 3). These results indicate that, in females, behavioral susceptibility may result from neuroanatomical changes that extend beyond the cortico-limbic circuit, encompassing motor, sensory and association areas.

**Figure 2.**
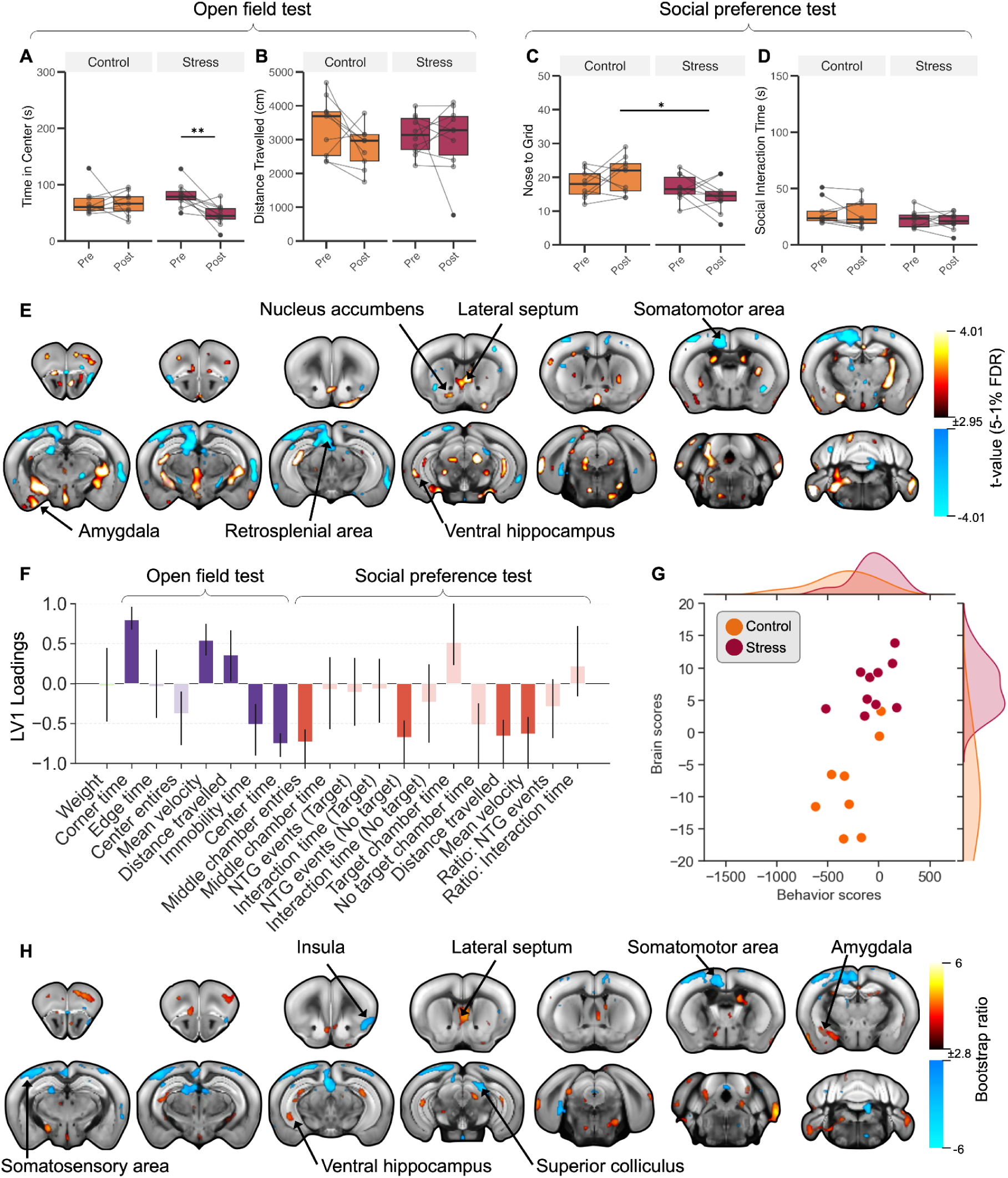
Covarying patterns of neuroanatomical and behavioral changes reveal female signature of stress susceptibility. Six days of CVS altered behavior in an open field test (A and B) and social preference test (C and D). (A) Stressed females spent less time in the center relative to baseline (p=0.0288, t-value=-2.29). In the social preference test, (C) stressed females showed decreased nose to grid events with a novel mouse (p=0.0114, t-value=2.027), without (D) significant changes in time in the social interaction zone. E) CVS led to significant volumetric changes in stressed females. Positive and negative t-statistics for volumetric brain changes: group by time-point interaction thresholded at 5-1% FDR, overlaid on the Allen CCFv3 template. Areas in warm colors (yellow-red) represent areas in which stress mice have more positive volumetric change relative to controls, whereas cool colors (blue) represent the opposite relationship. F-H) First significant LV of PLSC analysis on the difference across timepoints (p=0.039, covariance explained: 39.25%). F) Weights of each behavioral variable onto its respective LV, indicating the correlation of each behaviour to the pattern. Size of the bar is estimated through singular value decomposition, confidence intervals are calculated by bootstrapping. Confidence intervals that cross the zero line should not be considered as a significant contribution to the LV. Anxiety-like behavior in purple and social withdrawal in red. H) Weights of neuroanatomical changes onto its respective LV obtained from bootstrapping ratios overlaid on Allen CCFv3 template. Warm colors (yellow-orange) indicate positive relationships between volume change and behaviour, while cool colors (blue) represent negative relationships. G) Distribution of subject-specific brain and behavior scores, color coded by group. (NTG: Nose-to-grid)

### 2.2 Coordinated patterns of neuroanatomical changes associated with behavioral susceptibility in females

Six days of CVS exposure in females was sufficient to induce behavioural susceptibility and neuroanatomical changes that encompass limbic, association, motor and sensory areas. To determine whether these volumetric changes were associated with our signature of behavioral susceptibility, we used partial least squares correlation (PLSC) to uncover latent variables (LV) of maximal covariance between neuroanatomy and behavior, with each LV capturing an orthogonal phenotype of brain-behavior relationships.

We first estimated within-subjects behavioral and voxel-wise volume changes across time and examined their association using PLSC. This analysis revealed a significant LV (Figure 2F-H) that explained 39.25% of covariance and is characterized by increased volume in the ventral hippocampus, lateral septal complex, agranular insular cortex, nucleus accumbens and central and basolateral amygdala, alongside decreased volume in the somatosensory areas, retrosplenial area, anterior cingulate, mediodorsal nucleus of thalamus and superior colliculus (Figure 2H). This pattern of volumetric changes is associated with changes in both anxiety-and depressive-like behavior, specifically decreased time in center and increased locomotion in a novel environment, and decreased locomotion in a social environment (Figure 2F). This shared brain-behavior dimension was most strongly represented in stressed females as revealed by the subject-specific brain and behavior scores (Figure 2G). We then examined brain-behavior patterns specifically at the post-stress time-point in control and stress females. This identified a significant LV (Supplementary Figure 2), which explained 20.7% of the covariance, and reflected greater volume in the prelimbic, lateral septal complex, entorhinal area, medial amygdalar nucleus, and periaqueductal gray, as well as lower volume in the basolateral amygdalar nucleus, anterior cingulate area, retrosplenial area, and agranular insular area (Supplementary Figure 2B) that was uniquely associated with greater anxiety-like behavior (Supplementary Figure 2A). This brain-behavior phenotype was most strongly represented in stressed females, as revealed by the subject-specific brain and behavior scores (Supplementary Figure 2C). Together, these results suggest that anxiety-like behavior is associated with neuroanatomical changes in core stress-responsive regions, whereas depressive- and anxiety-like behavior encompass more widespread changes in cortical, limbic, association and sensory regions.

### 2.3 Network architecture of stress susceptibility signature in females

PLSC analysis identified coordinated patterns of volumetric changes associated with behavioral susceptibility in female mice. To determine putative regional targets for therapies that ameliorate this phenotype, we asked if these emergent patterns of change were part of a structural network synaptically-connected together by hub regions. First, we thresholded and binarized our PLS brain patterns at BSR ≤ ±2.57 (corresponding to p<0.01) and defined nodes as spatially continuous clusters of connected voxels. Directed connectivity between nodes was inferred from viral tract-tracing experiments from the Allen Brain Institute (Knox et al., 2019) by quantifying the overlap in tracer-defined connectivity profiles between pairs of nodes, followed by assessing statistical significance against a null distribution of spatially matched random connections (Supplementary Figure 1).

Using the Louvain algorithm (Lancichinetti & Fortunato, 2012), we evaluated community structure across a range of resolution parameters (0.3 ≤ 𝛾 ≤ 3) and quantified similarity between partitions using adjusted mutual information. A stable community partition was selected when the similarity across partitions was stable across a continuous resolution interval (Δ𝛾 ≥ 0.2). We identified 7 interconnected communities underlying the brain pattern associated with both depressive- and anxiety-like behavior previously described (Figure 3). The first community encompassed sensory areas, as well as the insula, prelimbic, dorsal striatum, entorhinal area and hippocampus. The second community encompassed mostly limbic structures, including the anterior cingulate area, hippocampus, lateral septal complex, and striatum. The third community encompassed regions associated with defensive behaviors (Lee et al., 2022), including the periaqueductal gray, retrosplenial area and superior colliculus. The fourth and fifth communities encompassed primarily cerebellar and midbrain regions. The spatial distribution of the sixth community recapitulated that of the salience network in rodents (Gozzi & Schwarz, 2016; Mandino et al., 2021), including regions like the insula, amygdala, hippocampus and striatum. Finally, the seventh community recapitulated the spatial distribution of the default-mode network (Gozzi & Schwarz, 2016), including regions like the anterior cingulate area, retrosplenial area and thalamus.

**Figure 3.**
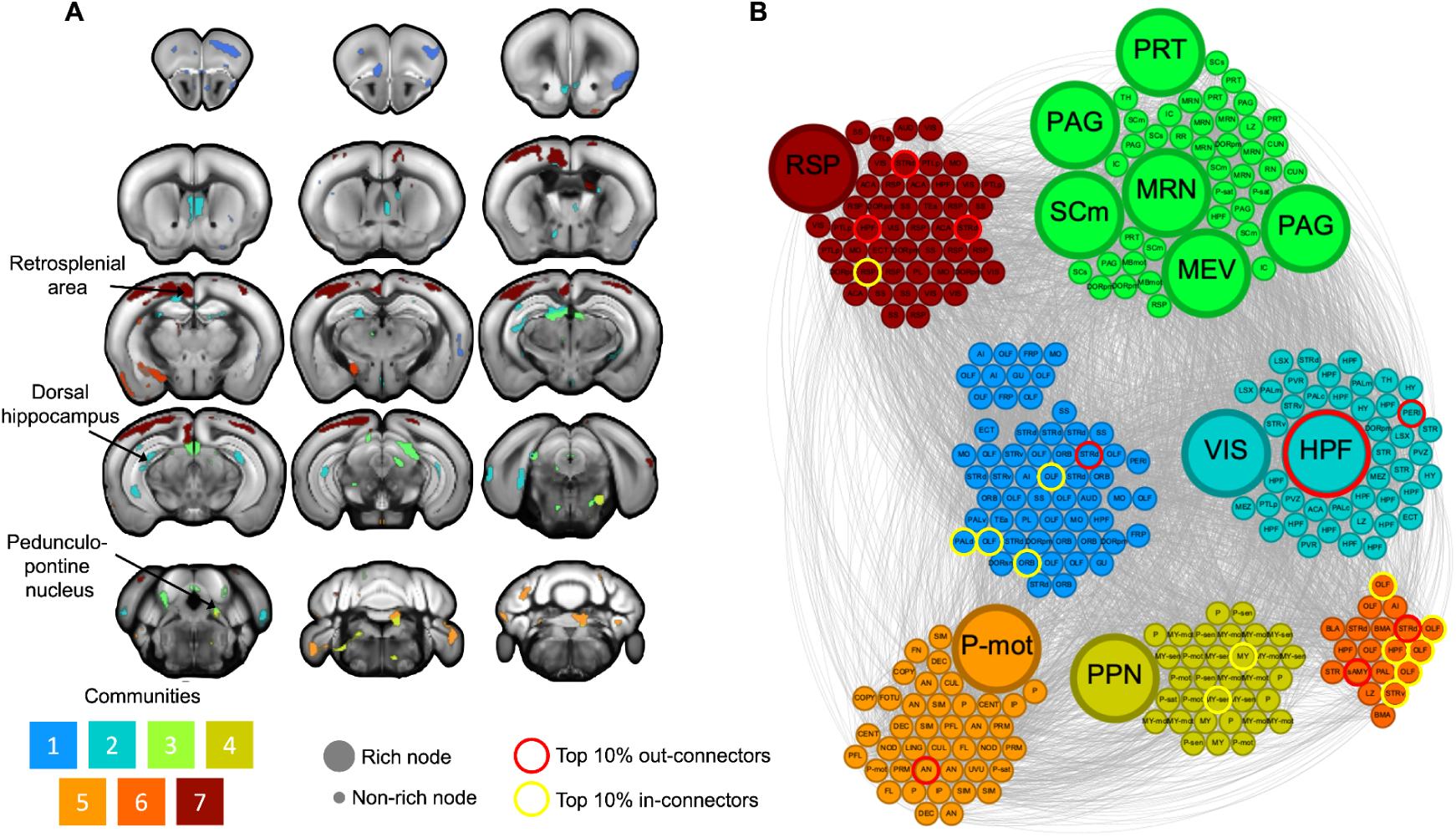
Network structure of significantly connected voxels from shared brain-behavior dimension in females. This LV captured brain changes across time that are associated with increased anxiety-like behavior and changes in locomotion in females most prominently in stressed female mice A) Color of each voxel in the coronal brain slices denotes community membership. B) Graphic representations of each network, the color of the nodes denotes community membership, and size represents whether the node is part of the rich club. The acronyms in each node denote the brain region where it is found. (RSP: retrosplenial area; PAG: periaqueductal gray; SCm: superior colliculus, motor related; MRN: midbrain reticular nucleus; MEV: midbrain trigeminal nucleus; VIS: visual areas; HPF: hippocampal formation; P-mot: pons, motor related; PPN: Penduculopontine nucleus. Label acronyms for non-rich nodes are described in Supplementary Table 1).

In the connectome, the most highly connected nodes, termed ‘rich club nodes’, are thought to be especially important in integrating information and by virtue of their many connections, they can make the network resilient to disruptions. Their high connectedness also predicts that alterations in these nodes will have large impacts on network function. To define rich club membership, we first quantified the degree (total number of connections) of each node and computed the normalized rich club coefficient (Fulcher & Fornito, 2016), defined as the ratio of the empirical rich-club coefficient and the mean rich club coefficient of 1000 rewired networks. The topological rich club regime was defined as the range of degrees for which the normalized rich-club coefficient was statistically significant against this null distribution (Supplementary Figure 5). Nodes were classified as part of the rich club if their degree was greater or equal to the mean plus one standard deviation of this range (Fulcher & Fornito, 2016).

In this brain pattern, the rich club consisted of nodes associated with visual processing (the superior colliculus and visual regions), locomotion (the pedunculopontine nucleus) (Garcia-Rill et al., 2019), food intake (the midbrain trigeminal nucleus) (Fortin et al., 2021), social and threat behaviors (the periaqueductal gray) and cortical regions including the hippocampus and retrosplenial area (Figure 3B).

These results suggest that in females, stress induced coordinated neuroanatomical changes across interconnected brain circuits, and that alterations to hub regions that compromise efficient communication between communities may contribute to behavioral susceptibility.

### 2.4 Spatial gene expression analyses reveal potential biological mechanisms of female susceptibility

Analyzing neuroanatomical and behavior changes in females revealed stress susceptibility signatures in which coordinated volumetric changes across key brain regions involved in the communication between interconnected communities associated with behavioral susceptibility. We then considered potential biological mechanisms underlying this brain-behaviour dimension of stress-susceptibility, focusing on the role of regional variation in gene expression. We used the Allen Institute for Brain Science’s gene expression dataset in C57BL/6 mice (Lein et al., 2006) to identify genes associated with the neuroanatomically defined stress-susceptibility signatures. We separately considered associations to positive or negative volume changes. Calculating the spatial correlation coefficient between the female within-subjects LV1 and the normalized gene expression for each of the 4073 available genes yielded a number of genes that were associated with positive volume change. The top correlated genes included Chrm3, Gtdc1, Khdrbs3, Zbtb16 and Satb1, and were enriched for the GO term of “regulation of protein localization to cell surface”, suggesting that epigenetic mechanisms may lead to alterations in cholinergic signaling (Matsui et al., 2000; Picciotto et al., 2012). Examining correlations with negative volume change identified a number of genes but these were not enriched for any GO terms.

### 2.5 Behavioral and neuroanatomical changes of male susceptibility

Having uncovered neuroanatomical signatures of behavioral susceptibility in females, we then used a phenotype-matched design to examine behavioral and neuroanatomical changes induced by 28 days of CVS in male mice (Johnson et al., 2021; Muir et al., 2020). Linear mixed effects models revealed that, similar to 6 days CVS in females, in males, 28 days of CVS increased anxiety-like behavior as indicated by decreased time spent in the center of the open field (Figure 4A). However in males, CVS did not significantly alter measures of social behavior (Figure 4C and D).

**Figure 4.**
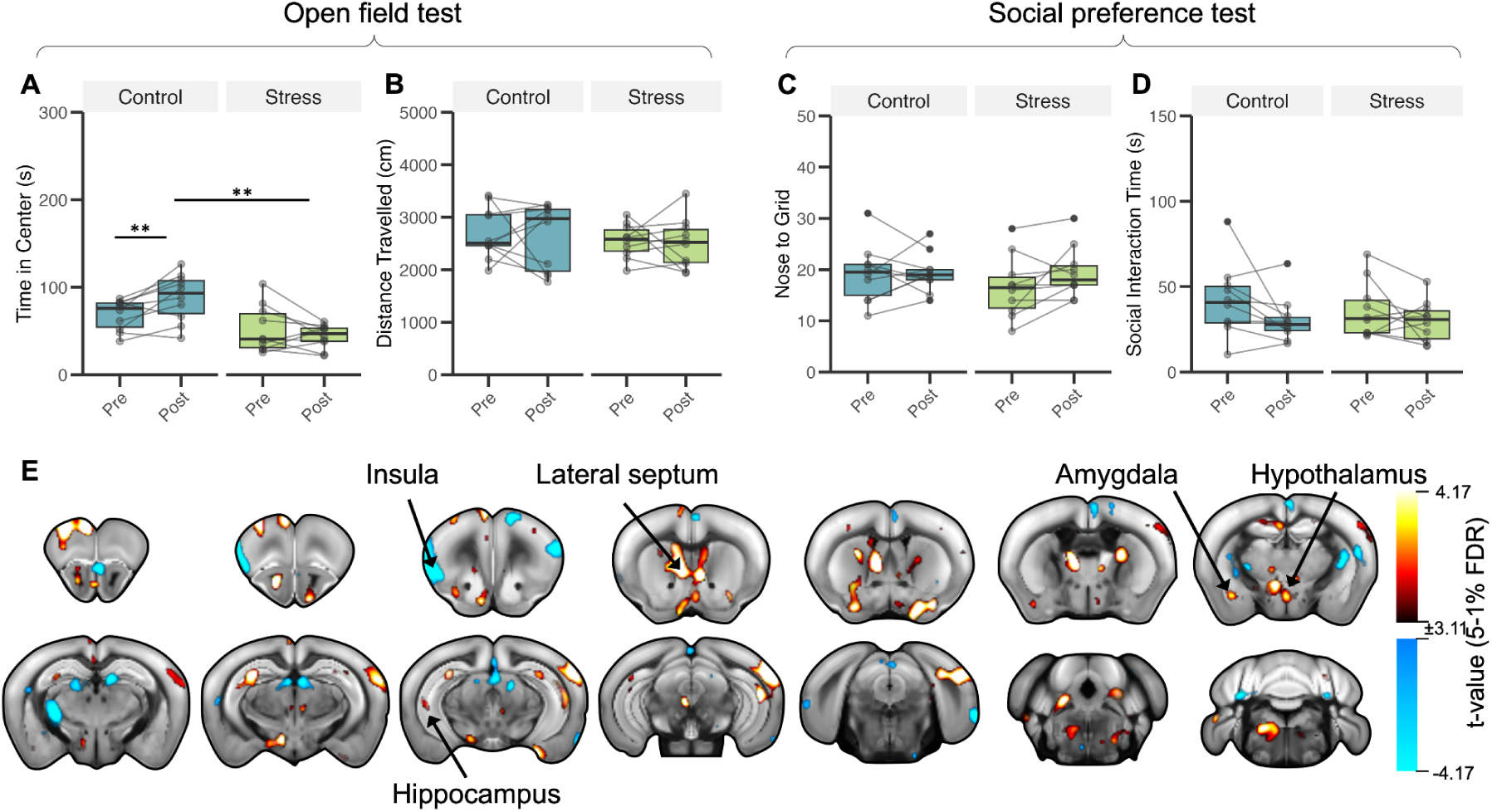
Behavioral and neuroanatomical changes observed after 28 days of CVS in males. (A) Stressed males spent less time in the center relative to controls (p=0.0006, t-value=4.522), without (C and D) significant differences in the social preference test. E) Significant volumetric changes in stressed males compared to controls. Positive and negative t-statistics for volumetric brain changes: group by time-point interaction thresholded at 5-1% FDR, overlaid on the Allen CCFv3 template. Areas in warm colors (yellow-red) represent areas in which stress mice have more positive volumetric change relative to controls, whereas cool colors (blue) represent the opposite relationship.

As in female mice, we then used deformation-based morphometry to investigate voxel-wise volumetric changes derived from T1w structural MRI scans. In males exposed to 28 days of CVS, we observed the greatest differential volume increases in the lateral septal complex, temporal association areas, central amygdala, nucleus reuniens, nucleus accumbens and ventral hippocampus; and the largest differential volume decreases in the agranular insular cortex, somatomotor areas, gustatory areas, superior colliculus and simple lobule of the cerebellum (Figure 4E and Supplementary Figure 4).

As stress duration was titrated to induce comparable behavioral adaptations in males and females, a direct statistical comparison between sexes was not appropriate (Makin & Orban de Xivry, 2019). Therefore, we limited our evaluation of shared and “sex-specific” (referring to effects observed qualitatively in only one sex) neuroanatomical changes to a qualitative comparison relative to the above reported findings in females (Figure 5). As in our analysis in females, we observed volume increases in key stress response and mood regulation regions, including the nucleus accumbens, amygdala, lateral septal nucleus, prelimbic area, periaqueductal gray and medial preoptic area, alongside volume decreases in cortical regions like the insula, and somatomotor, anterior cingulate and retrosplenial areas. Regions showing opposite direction of change encompassed areas that process and integrate sensory and interoceptive information, and relay it to limbic and cortical regions. These included temporal association, visceral visual and auditory areas, nucleus reuniens, mediodorsal nucleus of the thalamus and superior colliculus. Lastly, a subset of regions, such as the lateral amygdalar nucleus, paraventricular nucleus of the hypothalamus and medial habenula, exhibited changes exclusively in males; while regions including the perirhinal area, paraventricular nucleus of the thalamus and red nucleus were only altered in females. Together these findings suggest that while the state of stress susceptibility involves similar changes in core limbic regions across both sexes, association and hypothalamic regions may undergo sex-specific remodelling.

**Figure 5.**
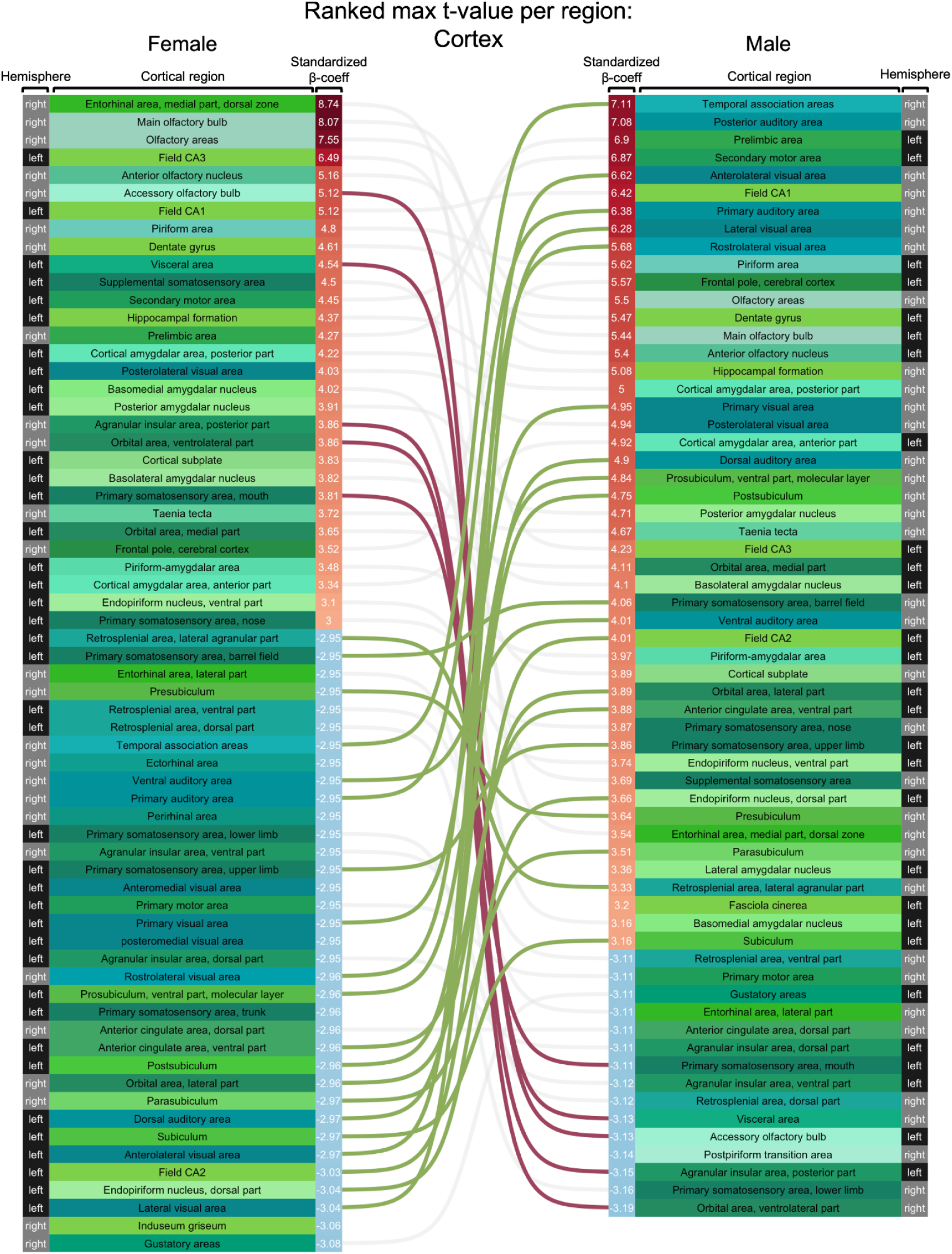
Phenotype-matched durations of CVS induce volume changes in shared and “sex-specific” regions in the cortex. Ranked region-wise changes in volume for females (left) and males (right). The red and blue shading of □-coefficient values denote volume increase or decrease, respectively, and the values denote the maximum standardized beta-coefficient found in each region extracted from the results of the voxelwise linear mixed effects models shown in Figures 2 and 4. Lines connect similar brain regions with significant volume change in each sex, pink lines denote regions with positive volume change in females and negative change in males, green lines denote regions with positive volume change in males and negative change in females and gray lines denote regions with similar direction of change in both sexes.

### 2.6 Coordinated patterns of neuroanatomical changes associated with behavioral susceptibility in male

We then used PLSC to examine how the identified neuroanatomical changes associate with behavioral susceptibility. We did not identify any significant LVs capturing the association between within-subject measures of changes in brain volume and behavior across time as we had observed in females, however we identified two significant LVs capturing brain-behavior relationships at the post-stress time-point. LV1 explained 29.4% of the covariance (Figure 6A-C) and is characterized by greater volume in the cortical amygdala, visual areas, nucleus accumbens, hypothalamus, lateral habenula and mediodorsal nucleus of the thalamus, and lower volume in the entorhinal area. These volume alterations were associated only with changes in depressive-like behavior, including greater social preference as well as lower body weight, and were most strongly represented in stressed males as indicated by the subject-specific scores. LV2 (Figure 4D-F), which explained 27.06% of the covariance, captured greater volume in the basolateral amygdala, hippocampus, lateral septal complex and auditory areas, and lower volume in the infralimbic area, agranular insular cortex and gustatory areas. These volume alterations were associated with changes in anxiety-like behavior, including decreased time in center and increased locomotion, and were most strongly represented in stressed males, as indicated by the subject-specific brain and behavior scores.

**Figure 6.**
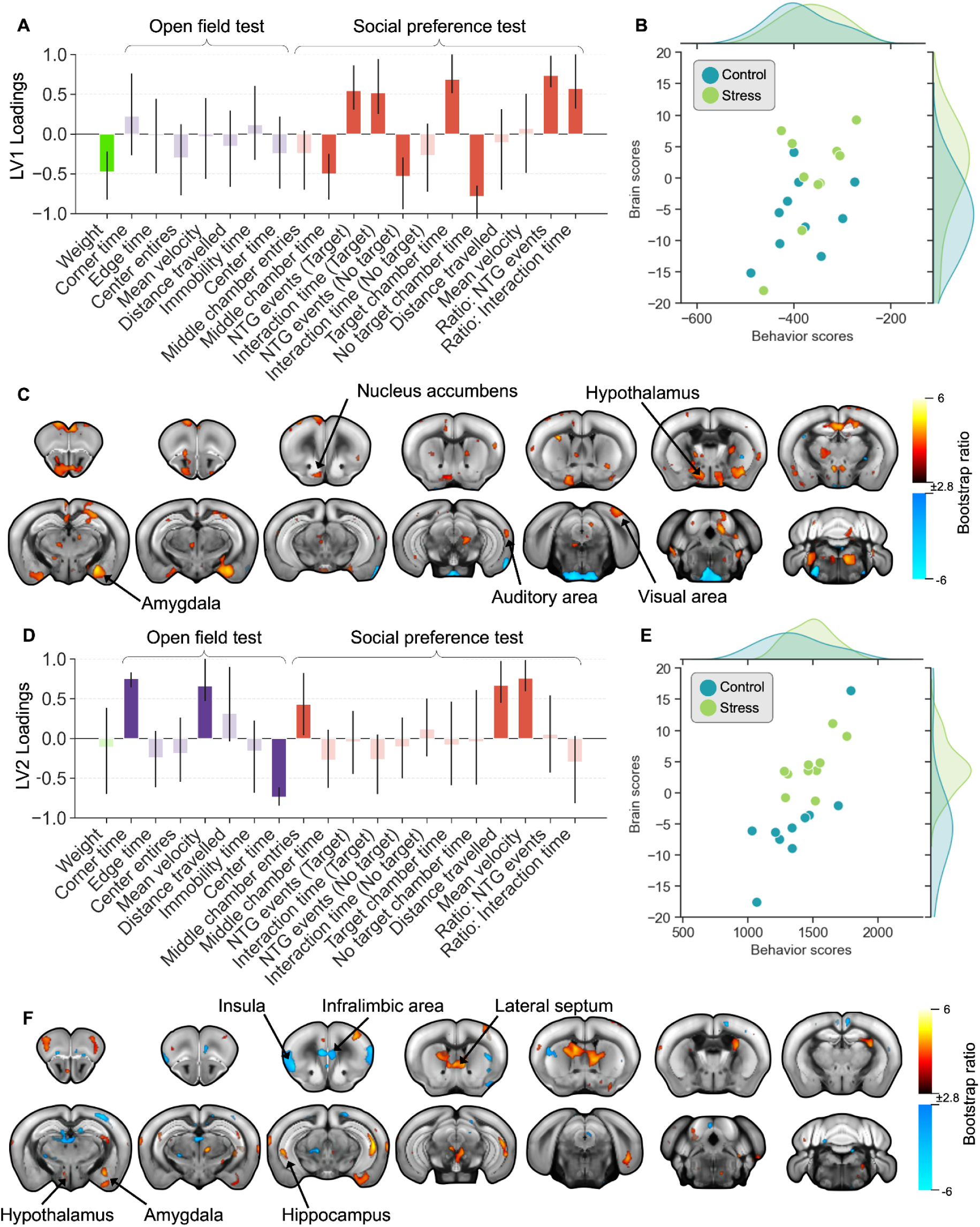
Covarying patterns of neuroanatomical and behavioral changes reveal two male signatures of stress susceptibility. A-C) LV1 of PLSC analysis on post-CVS measures (p=0.032, covariance explained: 29.40%). D-F) LV2 of PLSC analysis on post-CVS measures (p=0.013, covariance explained: 27.06%). A and D) Weights of each behavioral variable onto its respective LV, indicating the correlation of each behaviour to the pattern. Size of the bar is estimated through singular value decomposition, confidence intervals are calculated by bootstrapping. Confidence intervals that cross the zero line should not be considered as a significant contribution to the LV. Anxiety-like behavior in purple and social withdrawal in red. C and F) Weights of neuroanatomical changes onto its respective LV obtained from bootstrapping ratios overlaid on Allen CCFv3 template. Warm colors (yellow-orange) indicate positive relationships between volume change and behaviour, while cool colors (blue) represent negative relationships. B and E) Distribution of subject-specific brain and behavior scores, color coded by group. (NTG: Nose-to-grid)

These results suggest that, unlike females where we identified a shared signature of stress susceptibility, in males stress induced two potentially orthogonal processes: one associated with anxiety-like behavior and another with changes in social preference, each reflecting distinct patterns of brain alterations. Although we observed that several regions exhibited similar neuroanatomical remodeling across sexes, intriguingly their association to behavior differed between sexes. This dissociation suggests that similar neuroanatomical changes may mediate different behavioral adaptations in males and females, potentially reflecting sexually differentiated roles of these regions within distributed networks.

### 2.7 Network architecture and potential biological processes of stress susceptibility signatures in males

In contrast to females, where we observe a single dimension of volumetric alterations associated with anxiety- and depressive-like behaviors, our PLSC analysis in males revealed two orthogonal dimensions of volumetric alterations associated with depressive- or anxiety-like behavior. We then examined the topology of the structural network underlying these brain-behavior correlates to investigate the role of these alterations within a network framework. In the connectome corresponding to the depressive-like brain-behavior dimension, we identified 8 communities (Figure 7A and B). These included: (1) a community encompassing thalamic, somatosensory, striatal and retrosplenial area; (2) a hippocampal community; (3) a cerebellar community; (4) a frontal and thalamic community; (5) a subcortical community that included the amygdala, striatum, insula and pallidum; (6) a medullary community; (7) a brainstem community that included the periaqueductal gray and reticular nuclei; and (8) a hypothalamic community. Here, the rich club encompassed hypothalamic, brainstem, and amygdalar regions.

**Figure 7.**
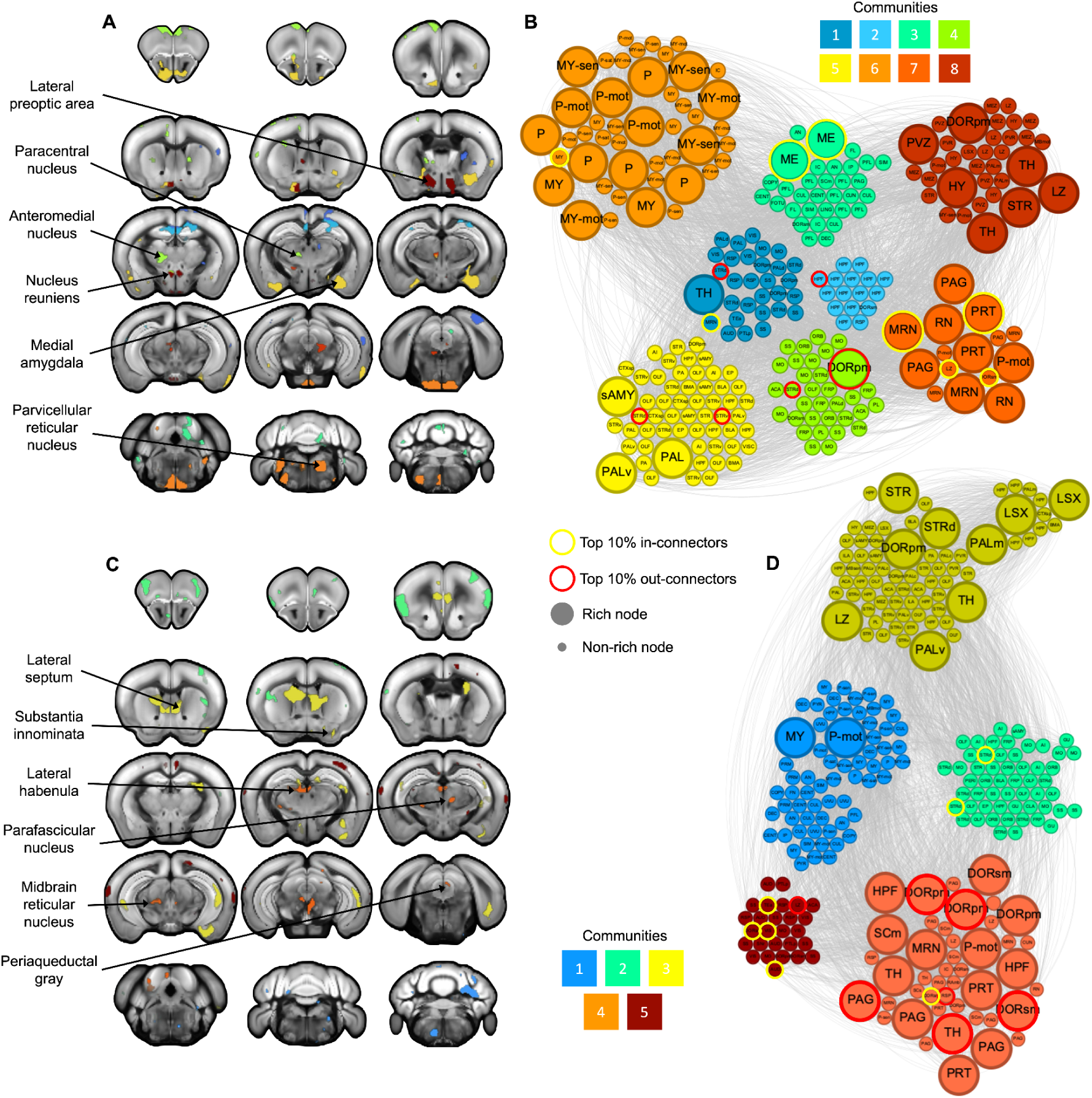
Network structure of significantly connected voxels from post-stress LVs in males. A and C) Color of each voxel in the coronal brain slices denotes community membership. B and D) Graphic representations of each network, the color of the nodes denotes community membership, and size represents whether the node is part of the rich club. The acronyms in each node denote the brain region where it is found. (MY-sen: medulla, sensory related; P-mot pons, motor related; P: pons; ME: median eminence; TH: thalamus; sAMY: striatum-like amygdalar nuclei; PALv: pallidum ventral part; PAL: pallidum; DORpm: thalamus, polymodal association cortex related; DORsm: thalamus, sensory-motor cortex related; MRN: midbrain reticular nucleus; PRT: pretectal regions; PAG: periaqueductal gray; RN: red nucleus; HY: hypothalamus; PVZ: periventricular zone; STR: striatum; LZ: hypothalamic lateral zone; LSX: lateral septal complex; PALm: pallidum, medial region; HPF: hippocampal formation; SCm: superior colliculus, motor related; Label acronyms for non-rich nodes are described in Supplementary Table 1).

In the connectome underlying the brain pattern captured by the anxiety-like brain-behavior dimension, we identified 5 communities (Figure 7C and D). These included: (1) a community that mostly encompassed cerebellar and medullary regions; (2) a community similar to the sixth community in females whose spatial distribution recapitulated that of the salience network in rodents (Mandino et al., 2021), including regions like the insula, amygdala, hippocampus and striatum; (3) a limbic community that encompassed the anterior cingulate area, amygdala, hippocampus, infralimbic area, lateral septal complex and striatum; (4) a community similar to the third community in females, which encompassed regions associated with defensive behaviors (Lee et al., 2022), including the periaqueductal gray, thalamus and superior colliculus; and (5) a community similar to the seventh community in females, which recapitulated the spatial distribution of the default-mode network (Gozzi & Schwarz, 2016), including regions like the anterior cingulate area, retrosplenial and visual areas. Here, the rich-club encompassed limbic, hippocampal and thalamic nodes.

Furthermore, while spatial gene expression analyses in males (both post-stress LV1 and LV2) identified genes correlated with positive and negative volume differences, these were not enriched for any GO terms.

Together, these results suggest that in males, distinct patterns of neuroanatomical remodelling associated with depressive- or anxiety-like behavior implicate segregated structural networks, suggesting a modular organization supporting distinct behavioral domains. In contrast, females exhibit a more integrated network architecture, in which a shared structural signature is associated with both behavioral domains. The greater number of rich club nodes observed in males may indicate greater network redundancy, potentially increasing its resilience to perturbations. However, in females, stress may preferentially target a smaller set of rich club nodes, whose alterations may propagate more easily across the network, affecting several behavioral domains simultaneously. Such qualitative differences in the topology of these networks may help explain why males require greater durations of stress exposure to reach behavioral susceptibility, whereas in females, perturbation of key nodes may more readily produce behavioral alterations.

## 3. Discussion

As the prevalence of depressive and anxiety disorders continues to increase, so too does the imperative to develop effective treatment and prevention strategies. To this end, including sex as a biological variable in preclinical investigations is essential given the pronounced sex differences in mood disorders (Seney & Sibille, 2014). Preclinical evidence investigating mechanisms of stress susceptibility in each sex suggest that similar behavioral alterations observed in both males and females can be attributed to both shared and distinct neurobiological mechanisms (Muir et al., 2020). However, the lack of phenotype-matched protocols, inclusion of females and brain-wide investigations have limited our understanding of the neurobiological alterations underlying stress-susceptibility in both sexes. Using phenotype-matched stress protocols, we developed the first brain-wide characterization of stress susceptibility signatures in male and female mice. Similar to our previous work (Muir et al., 2020), we showed that titrating the duration of CVS in a sex-specific manner results in comparable behavioral adaptations across both sexes. However, while in both sexes, stress susceptibility implicated comparable neuroanatomical changes, we observed that the direction of change, association to behavior and hub regions were distinct in each sex.

By examining neuroanatomical changes at the whole-brain level, we identified that stress-susceptibility involves neuroanatomical alterations in comparable brain regions between sexes. We show that regions identified by previous studies such as the nucleus accumbens, amygdala, hippocampus, hypothalamus, and prefrontal cortex (Bittar et al., 2021; Hodes et al., 2015; Liu et al., 2018; Muir et al., 2020) are involved in stress-susceptibility in both sexes. Previously uncharacterized regions in this model included the somatosensory cortex, agranular insular area, entorhinal area, visual areas, auditory areas and anterior cingulate area. Regions like the insula and the anterior cingulate area are part of the salience network, a network that has been shown to be enlarged in individuals with depression and is involved in interoceptive awareness (Dosenbach et al., 2025). The impaired sensory processing induced by alterations in these regions is suggested to underlie somatic symptoms observed in depression, including sleep disturbances, changes in appetite and pain symptoms (Chen et al., 2022; Kong et al., 2022; McGirr et al., 2020). Our whole-brain approach allowed for the unbiased discovery of previously unidentified regions that undergo stress-induced alterations and could suggest that CVS induces alterations in behavioral domains beyond depressive- and anxiety-like behavior, which require further characterization.

Using a data-driven approach, we mapped individuals into lower dimensional spaces using PLSC and identified latent dimensions of neuroanatomical remodelling associated with behavioral susceptibility to stress. This multivariate framework allowed us to capture neurobiological signatures that are behaviorally relevant across multiple domains and, rather than assigning categorical labels, to embrace the heterogeneity of stress-induced adaptations by positioning each subject along a continuum of brain-behavior expression. In females, volumetric change in regions, including the insular cortex, anterior cingulate area, amygdala and sensorimotor regions, were jointly associated with changes in anxiety- and depressive-like behavior. This pattern suggests that CVS changes the longitudinal trajectory of neuroanatomical and behavioral change across time. Previous studies on the bidirectional relationship between emotional processing and altered physiological states suggest the insular cortex as a key modulator of anxiety-like states associated with interoceptive processing of visceral signals (Hsueh et al., 2023). In conjunction with the anterior cingulate area and the amygdala, these regions are suggested to modify behavior by altering the integration of somatic and emotional information ((Bud) Craig, 2009). This finding suggests that in females, the alterations induced by stress result in a unitary effect on both anxiety- and depressive-like behavior, which may potentially mirror the increased prevalence of comorbid anxiety observed in women (Cavanagh et al., 2016). Although a similar pattern was observed in males (Supplementary Figure 6), it did not reach significance, further underscoring the relevance of this relationship in females.

In contrast, in males, stress-susceptibility mapped onto two orthogonal brain-behavior LVs. The first LV was characterized by greater anxiety-like behavior and volumetric alterations in regions including the lateral septal complex, prefrontal cortex, ventral hippocampus, basolateral amygdala and insula, and the second LV was characterized by depressive-like behavior and volume alterations in the dorsal hippocampus, hypothalamus, cortical amygdala and nucleus accumbens. While only a few studies have characterized sex-differences in the structure and function of these regions, these studies suggest that sex-differences in the innervation of circuits of the lateral septal complex are likely to influence social behavior in a sex-specific manner (Bredewold et al., 2014). By integrating environmental information encoded by the activity of structures like the hypothalamus, supramammillary nucleus, amygdala and hippocampus, the lateral septum modulates generalized anxiety states, social avoidance or locomotion (Besnard & Leroy, 2022; Li et al., 2022; Wirtshafter & Wilson, 2019, 2021). In contrast to females where stress susceptibility was captured by a single latent dimension, our results suggest that in males stress-induced adaptations in depressive- or anxiety-like behavior may be to be modulated by distinct neural substrates. Future work should examine whether this compartmentalized organization contributes to the greater resilience observed in males to shorter stress durations.

By leveraging the high-resolution and whole-brain coverage of MRI and interpreting our imaging results in the context of openly available datasets on neuronal connectivity and gene expression, we uncovered the roles of different regions implicated in each LV within each sex. We identified distinct normally interconnected communities that undergo neuroanatomical remodelling following CVS and are associated with emotional, sensory and social functions. Importantly, in both sexes we identified two communities whose spatial distribution recapitulated that of the salience and default-mode network (Gozzi & Schwarz, 2016; Mandino et al., 2021). In females, cortical regions like the retrosplenial and visual areas are key for communication, whereas in males, subcortical regions like the thalamus play an essential role. In addition to the evidence on sex-specific alterations in functional connectivity of these networks in depression (Yang et al., 2024), our results underscore the importance of characterizing the potential sex-differences that cause stress to have unitary vs orthogonal effects in females and males. Elucidating the role of these hub regions in each sex may lead to the development of targeted treatment strategies, like deep brain stimulation (Irmen et al., 2020), for females and males.

Lastly, our spatial gene expression analysis revealed genes whose spatial expression pattern is correlated to the brain pattern derived from PLSC. In particular, in females, regions whose volumetric changes were associated with depressive- and anxiety-like behaviors highly express genes previously implicated in the process of protein localization to the cell surface. The top correlated genes associated with this phenotype suggest that alterations observed in this LV may be associated with parasympathetic cholinergic signaling [Chrm3] (Matsui et al., 2000; Picciotto et al., 2012), as well as epigenetic regulators of cardiovascular functions [Zbtb16 and Miat] (Stilz et al., 2026). Studies in humans show that depression alters autonomic function differently in men and women, with depressed women exhibiting a greater risk for adverse cardiovascular events (Tobaldini et al., 2020). Future work examining the role of autonomic adaptations following chronic stress may elucidate critical mechanisms into the pathophysiology of depressive-like phenotypes in females.

As a field, we face a significant conundrum in basic neuroscientific research. We seek to examine the detailed architecture of brain signatures that underlie phenotypic expression yet our characterizations will remain limited until the historical exclusion of females across toolkits, methodologies and datasets is rectified. In spite of these limitations, we propose the analyses that we have shown as a means to best characterize the experimental data we present here. It should be noted that others have taken the same approach (Guma et al., 2024), but the impact that the male-bias has on our connectomic and transcriptomic results has yet to be characterized and fully understood. Evidence suggests that sexual differentiation in the rodent brain, either from sex-differences in genetics or gonadal hormones, takes place in the peri-and post-natal period (Premachandran et al., 2020; Qiu et al., 2018) and, in humans, studies suggest different topological properties in the structural connectivity of men and women (Tunç et al., 2016). Lastly, the behavioral paradigms employed in this study were optimized for male mice (Lopez & Bagot, 2021). The continued characterization of females is critical in a basic neuroscientific context to improve our understanding of health outcomes for the general population.

By employing a phenotype-matched approach, we provide the first whole-brain characterization of stress-susceptibility induced by CVS in female and male mice. Our findings underscore the importance of matching the experimental manipulation to capture the endophenotype of relevance across both sexes, even though it may require distinct manipulations for each sex. Future investigations using this type of approach to investigate mechanisms of stress susceptibility may elucidate targeted treatment strategies for males and females. Importantly, investigating how each sex responds to the same experimental manipulation provides complementary and valuable insights into understanding the role of sex differences in adaptations to chronic stress as a means to develop prevention strategies prior to disease onset.

## 4. Methods

### 4.1 Chronic Variable Stress (CVS)

Nine weeks-old female and male C57BL/6J mice were exposed to either 6 or 28 days of CVS (n=9F/10M control; n=10F/10M stress), as described in (Hodes et al., 2015). Mice were exposed to one of three 1 hour stressors daily: 100 random foot shocks (0.45 mA/1sec; administered in same-sex groups of 10 mice), tail suspension (with climb stoppers to avoid mice from climbing their tails (Can et al., 2012)), and restraint inside a 50 mL falcon tube (with holes for air circulation) (See Supplementary Methods Section 2 for more details).

### 4.2 Behavioral tests

We used open field and social preference tests at baseline and 24-hours after the last stressor to examine changes in anxiety- and depressive-like behavior (See Supplementary Methods Section 3 for more details).

### 4.3 Magnetic resonance imaging (MRI)

In vivo T1-weighted (T1w) structural images (100um^3^ voxels, TR: 20ms, TE: 4.5ms) were acquired with a 7T Bruker Biospec 70/30 MRI scanner equipped with a Bruker 1H Quadrature Transmit/receive MRI CryoProbe under a mixture of dexmedetomidine (0.25 mg/kg/h) and 1% isoflurane, in a 80% air and 20% oxygen mixture. T1w images were preprocessed and then analyzed using a deformation-based morphometry (DBM) pipeline (Chung et al., 2001; Germann et al., 2025) (https://github.com/CoBrALab/twolevel_ants_dbm). In brief, we used linear and nonlinear registration tools (Avants et al., 2011) to align images from all time-points for each subject and generate subject-specific templates. This step is particularly sensitive to the modelling of within subject change. The subject-specific templates were then used to generate a population-level template. Log-transformed Jacobian determinants were calculated from the deformation fields of the transformations that map each individual image to the average, resampled to the population-level template and blurred with Gaussian smoothing using ∼0.085 mm full width half-maximum kernel to better conform normative distribution assumptions for statistical testing (Chung et al., 2001). Voxels with positive values represent volume expansions relative to the average, while negative values represent contractions. We used the Jacobian determinants only from the nonlinear transformations (relative Jacobians) for statistical analyses as a means of removing residual global linear transformations (attributable to differences in total brain size, a major source of variability between mice (Lerch et al., 2012; Valiquette et al., 2023)). Throughout this manuscript, the term brain volume is used to refer to voxel-wise volume.

Images were visually inspected to assess quality control following the guidelines in: http://github.com/CoBrALab/documentation/wiki/Mouse-QC-Manual-(Structural) (See Supplementary Methods Section 4 for more details).

### 4.4 Statistical analysis

As in (Guma et al., 2021, 2023; Valiquette et al., 2023), linear mixed-effects models were used to investigate a group-by-timepoint interaction (subject ID as random intercepts) in relative Jacobians and behavioral tests, and were corrected for false discovery rate (FDR)(Benjamini & Hochberg, 1995). To investigate latent dimensions of brain-behavior relationships we used partial least squares correlation (PLSC) (Guma et al., 2021; McIntosh & Lobaugh, 2004; Zeighami et al., n.d.). We investigated patterns of covariance on (1) measures of change across time in brain volume and behavior within each subject, and (2) between brain volume and behavior at the final time-point. PLSC computations were bootstrapped 1000 times to obtain confidence intervals. All analyses were performed separately for each sex using pyls (https://github.com/rmarkello/pyls) in Python version 3.6.8. (See Supplementary Methods Section 5 for more details).

### 4.5 Structural connectivity

We used a graph theoretic framework to examine the topography of structural connections between regions significantly associated with behavior derived from PLSC. Brain weights were thresholded at values corresponding to a 95% confidence interval, binarized and labeled as groups of connected voxels (referred to as nodes). Using voxel-wise models of the mouse connectome (Knox et al., 2019), we calculated the overlap between the spatial connectivity (thresholded at 90th percentile) pattern of each source node to every other target node. We evaluated against a null distribution of random connections to remove non-significant connections, and binarized the adjacency matrix. As in (Fulcher & Fornito, 2016), we computed the normalized rich club coefficient as the ratio between the empirical rich-club coefficient and the mean rich-club coefficient of 1000 rewired networks (Rubinov & Sporns, 2010). Communities were derived using the Louvain algorithm (Coletta et al., 2020). To classify potential roles of our groups of voxels according to their intra- and inter-community connections, we calculated the participation coefficient (Guimerà & Nunes Amaral, 2005) where a participation coefficient closest to 1 reflects a higher distribution among all communities. Structural connectivity matrices were derived using Python version 3.8.2 and graph theory analyses were performed using the Brain Connectivity Toolbox (https://sites.google.com/site/bctnet/) on Matlab version 2020b. (See Supplementary Methods Section 6 for more details).

### 4.6 Spatial Gene Expression

To investigate potential molecular mechanisms underlying the neuroanatomical differences captured by the significant latent variables identified from our PLSC analyses, we sought to identify genes with most similar spatial expression patterns to these latent variables. We used the Allen Institute’s coronal mouse gene expression dataset in C57BL/6 male mice, which after processing consisted of 4071 normalized gene expression images (3836 unique brain-expressed genes). We computed the Spearman correlation between spatial gene expression patterns and the brain weights from PLSC, ranked these correlations, and identified gene ontology (GO) terms with genes that were preferentially distributed at the extreme ends of the ranked list (Ashburner et al., 2000; Merico et al., 2010). We quantified enrichment by calculating the area under the curve (AUC) formed by the cumulative distribution of genes within a module as a function of rank (Hoops et al., 2024). Statistical significance was evaluated against a null distribution of random phenotypes and corrected for FDR. Analyses were performed R version 3.5.1 (See Supplementary Methods Section 7 for more details).

## Acknowledgements

J.M. received support from the Canadian Institutes of Health Research (CIHR post-doctoral training award 202210MFE-491520-297096). We would like to thank Dr. Bruno Giros for lending us their learned helplessness apparatus.

---

**Figure 5 - Figure supplement 1.**
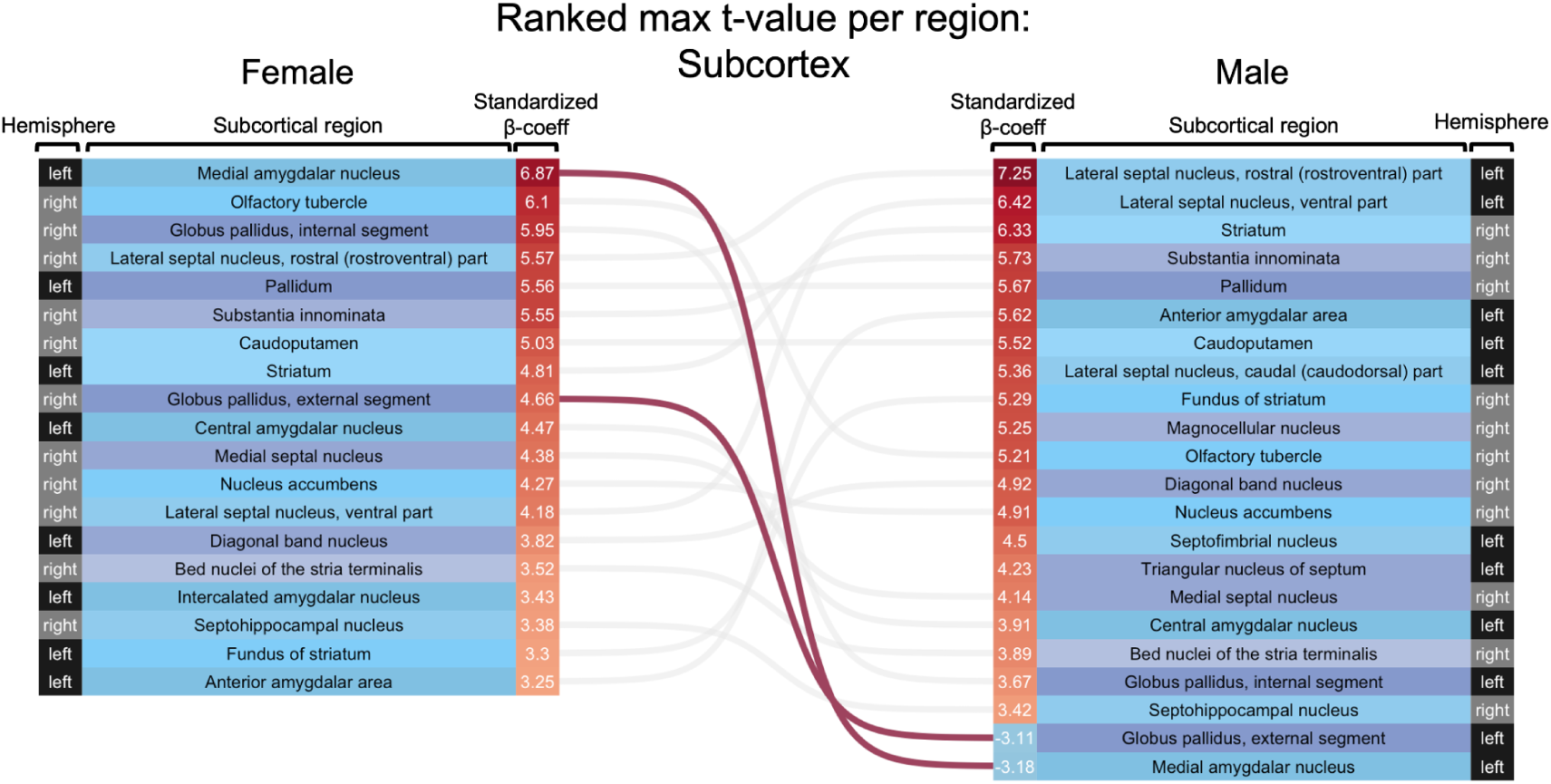
Phenotype-matched durations of CVS induce volume changes in shared and “sex-specific” regions in the cortex. Ranked region-wise changes in volume for females (left) and males (right). The colors red and blue denote volume increase or decrease, respectively, and the values denote the maximum standardized beta-coefficient found in each region extracted from the results of the voxelwise linear mixed effects models shown in Figures 2 and 4. Lines connect similar brain regions with significant volume change in each sex, pink lines denote regions with positive volume change in females and negative change in males, gray lines denote regions with similar direction of change in both sexes. No instances of positive volume change in males and negative change in females were observed.

**Figure 5 - Figure supplement 2.**
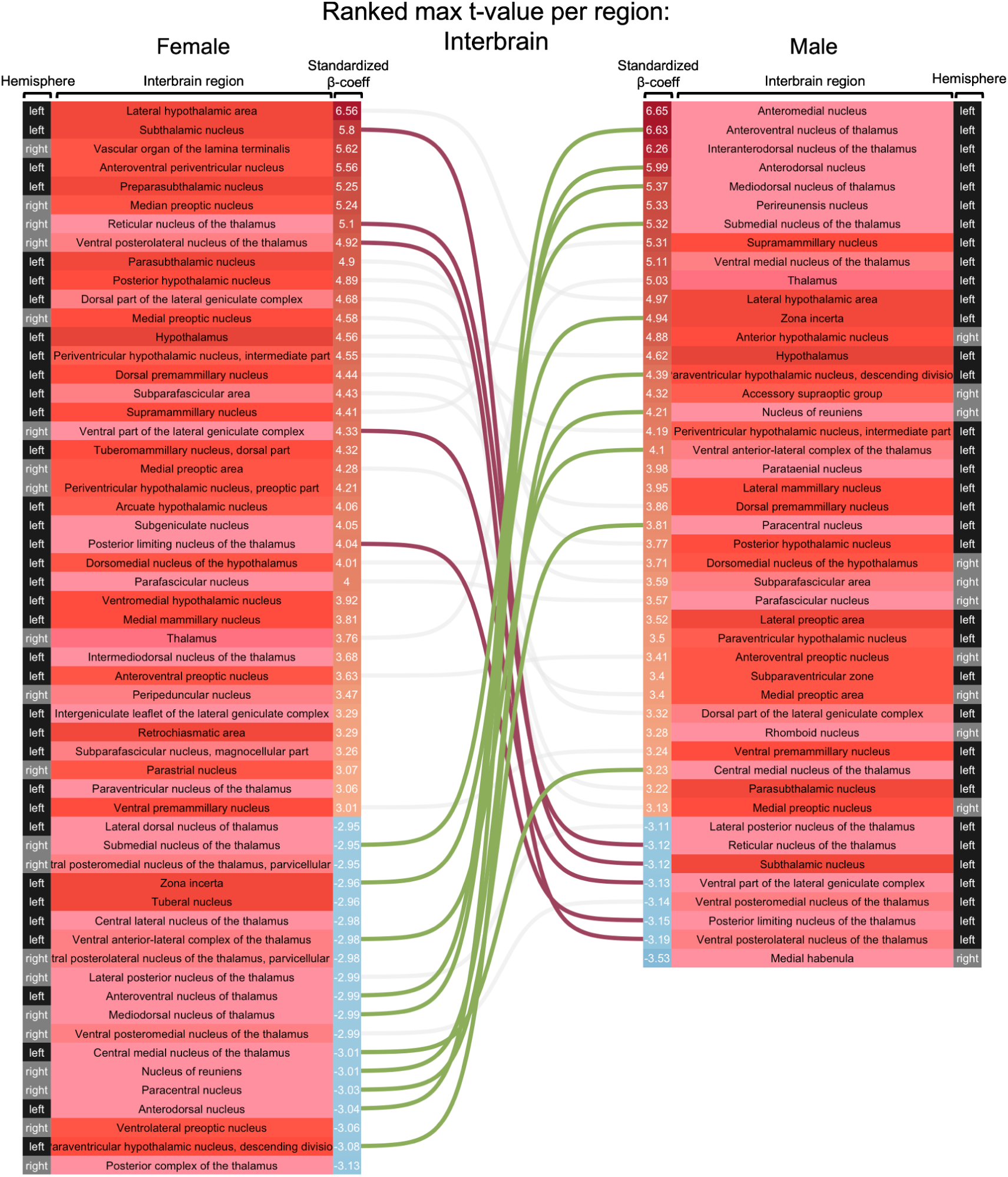
Phenotype-matched durations of CVS induce volume changes in shared and “sex-specific” regions in the interbrain. Ranked region-wise changes in volume for females (left) and males (right). The colors red and blue denote volume increase or decrease, respectively, and the values denote the maximum standardized beta-coefficient found in each region extracted from the results of the voxelwise linear mixed effects models shown in Figures 2 and 4. Lines connect similar brain regions with significant volume change in each sex, pink lines denote regions with positive volume change in females and negative change in males, green lines denote regions with positive volume change in males and negative change in females and gray lines denote regions with similar direction of change in both sexes.

**Figure 5 - Figure supplement 3.**
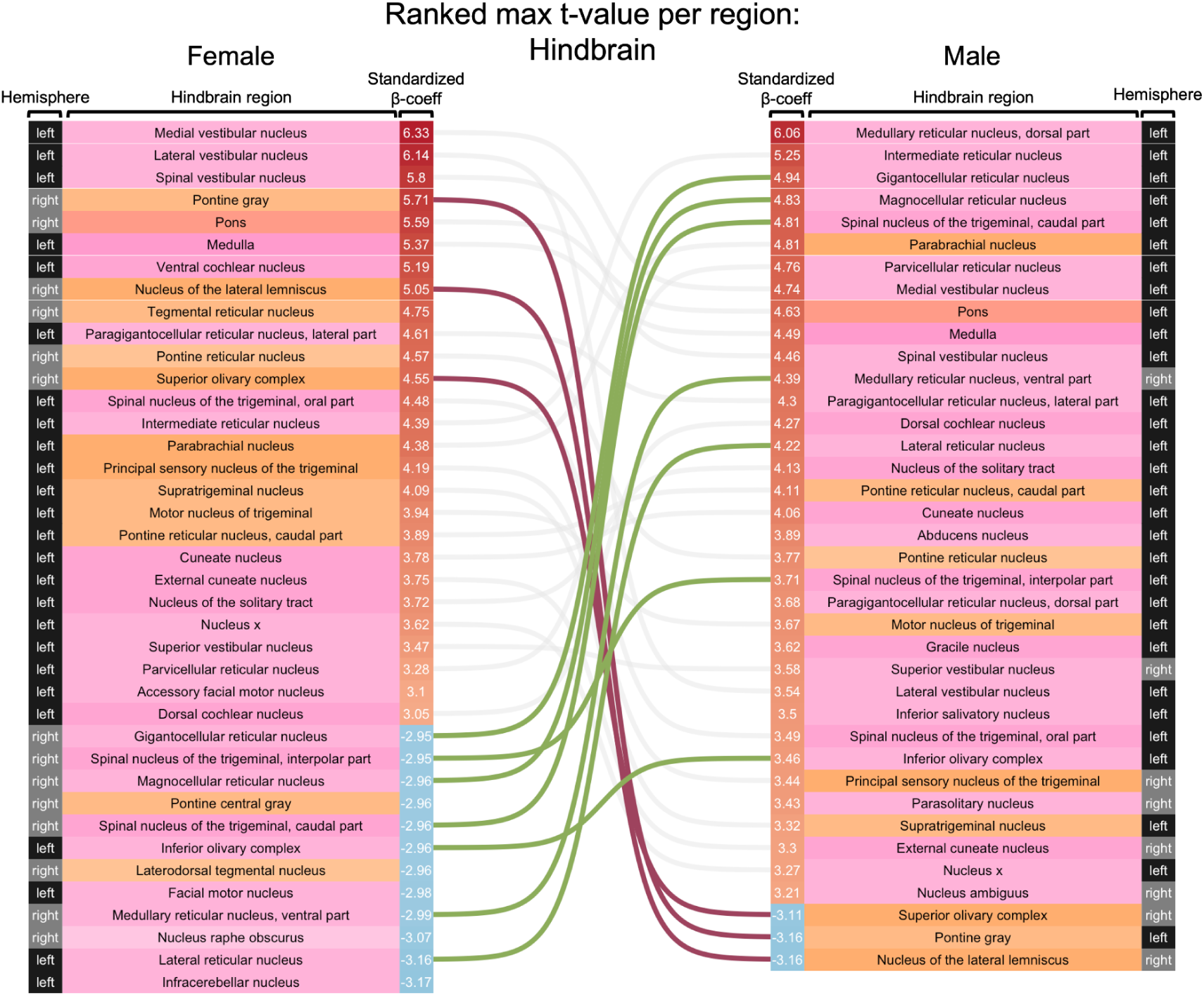
Phenotype-matched durations of CVS induce volume changes in shared and “sex-specific” regions in the hindbrain. Ranked region-wise changes in volume for females (left) and males (right). The colors red and blue denote volume increase or decrease, respectively, and the values denote the maximum standardized beta-coefficient found in each region extracted from the results of the voxelwise linear mixed effects models shown in Figures 2 and 4. Lines connect similar brain regions with significant volume change in each sex, pink lines denote regions with positive volume change in females and negative change in males, green lines denote regions with positive volume change in males and negative change in females and gray lines denote regions with similar direction of change in both sexes.

**Figure 5 - Figure supplement 3.**
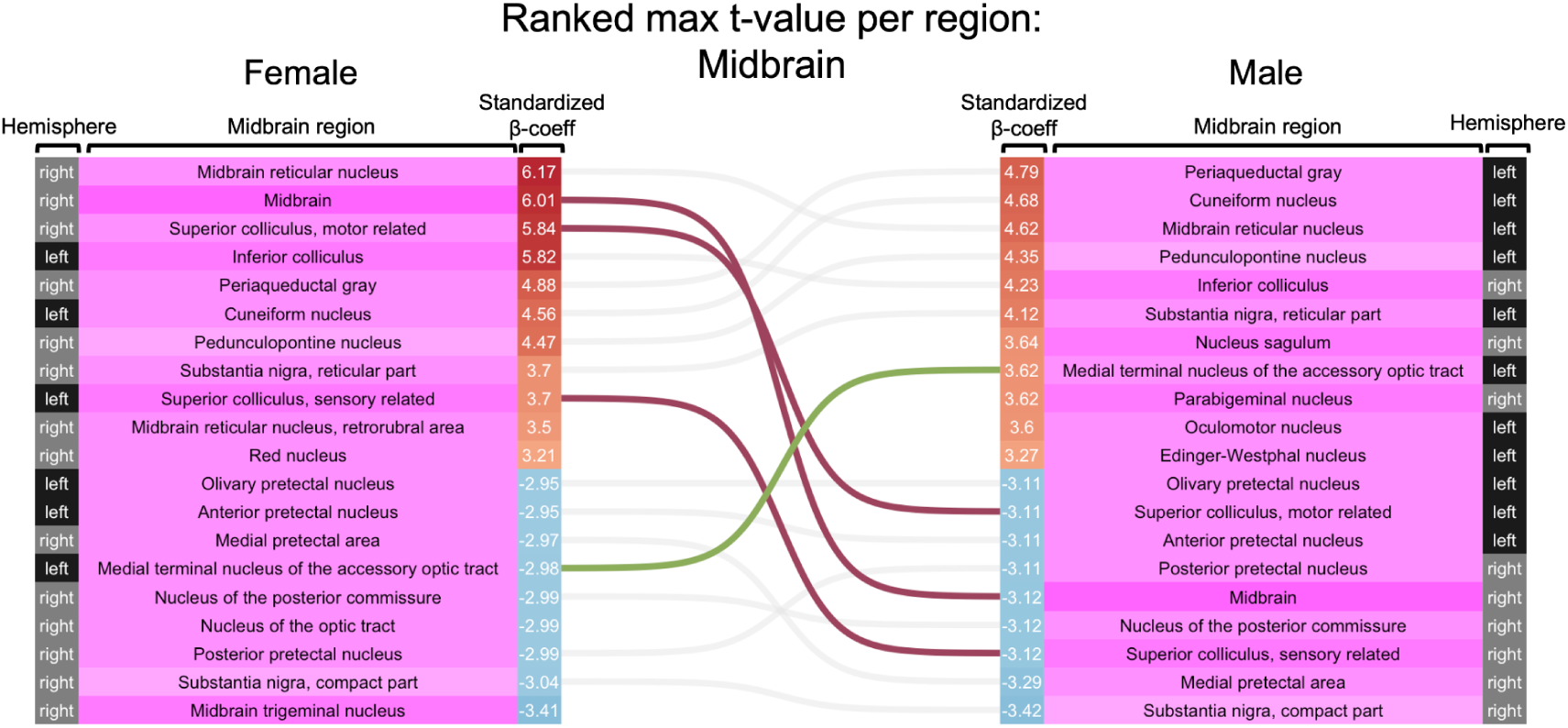
Phenotype-matched durations of CVS induce volume changes in shared and “sex-specific” regions in the midbrain. Ranked region-wise changes in volume for females (left) and males (right). The colors red and blue denote volume increase or decrease, respectively, and the values denote the maximum standardized beta-coefficient found in each region extracted from the results of the voxelwise linear mixed effects models shown in Figures 2 and 4. Lines connect similar brain regions with significant volume change in each sex, pink lines denote regions with positive volume change in females and negative change in males, green lines denote regions with positive volume change in males and negative change in females and gray lines denote regions with similar direction of change in both sexes.

**Figure 5 - Figure supplement 4.**
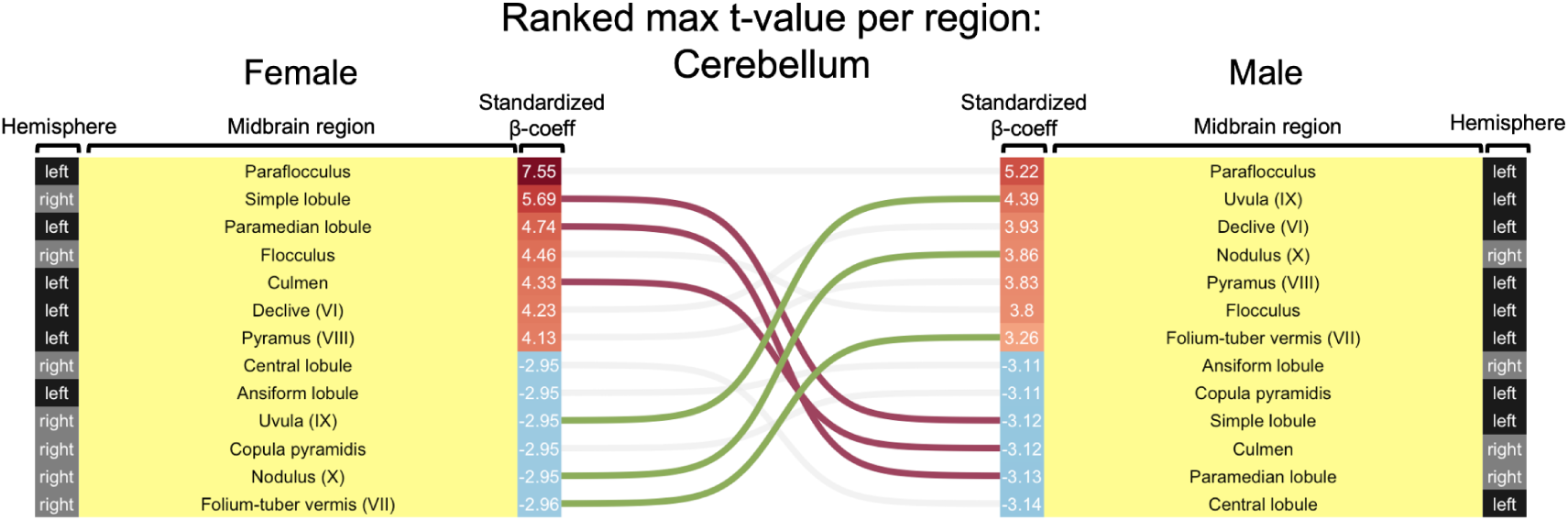
Phenotype-matched durations of CVS induce volume changes in shared and “sex-specific” regions in the cerebellum. Ranked region-wise changes in volume for females (left) and males (right). The colors red and blue denote volume increase or decrease, respectively, and the values denote the maximum standardized beta-coefficient found in each region extracted from the results of the voxelwise linear mixed effects models shown in Figures 2 and 4. Lines connect similar brain regions with significant volume change in each sex, pink lines denote regions with positive volume change in females and negative change in males, green lines denote regions with positive volume change in males and negative change in females and gray lines denote regions with similar direction of change in both sexes.

