## Supplementary Information for "Whole brain dimensional approach identifies networks of stress susceptibility in male and female mice"

**Supplementary Methods**

1. **Animals**

All procedures were approved by the McGill Animal Care Committee and were conducted in accordance with guidelines of McGill University's Comparative Medicine. 7-week-old male and female C57BL/6 mice were purchased from Jackson Laboratories and housed on a 12-hour light-dark cycle (lights on at 8:00 A.M. EST) at 20-25 C, group housed with 4-5 cage same-sex cage mates with ad libitum access to food and water for 10-15 days prior to the start of any manipulations. Mice (n=9F/10M control; n=10F/10M stress) were 9 weeks-old at the beginning of all experiments. All manipulations were performed during the light cycle following the experimental timeline in Figure 1A.

2. **Chronic variable stress**

Female and male mice were exposed to 6 or 28 days of CVS, respectively, as described in (Hodes et al., 2015). Mice were exposed to one of three stressors for 1 hour daily: 100 random foot shocks (0.45 mA/1sec; administered in same-sex groups of 10 mice), tail suspension (with climb stopper to avoid mice from climbing their tails (Can et al., 2012)), and restraint inside a 50-mL falcon tube (with holes for air circulation). On the first day, mice were habituated to the stress room prior to the first stressor. Stressors were administered at a variable time between 8am and 6pm to prevent habituation and in a different room from the behavioral testing room. For both sexes, stressors were presented in the same order across days. The number of days for CVS for males and females was determined following previous work from our laboratory (Muir et al., 2020) and based on previously identified sex differences in stress susceptibility (Hodes et al., 2015).

#### 3. Behavioral tests

Mice were habituated to the behavioral room 1 hour prior to testing. We conducted one test per day between 8AM and 2PM. Males and females were tested on different days. All equipment was cleaned with Prevail disinfectant cleaner between sessions. Recordings from the behavioral tests were analyzed with Ethovision XT 12 tracking system (Noldus Information Technology, Leesburg, VA, USA).

- **Open field test:** Mice were placed in the corner of an open gray plastic arena (45 cm x 45 cm) under red light and allowed to explore freely for 5 min. The following measures were recorded using Ethovision for statistical analyses: time in the center (40% of the arena: 27 cm x 27 cm central zone), frequency of passes through the center, time in edges, time in corners, total distance traveled, mean velocity and time immobile.
- **Social preference test:** Prior to testing, all mice (including novel conspecific social targets) were single housed overnight. This test consisted of two stages, each lasting 5-min under red light. During the first-stage (habituation stage), mice were placed in a three-chamber gray plastic box (26 (l) x 21.6 (w) x 21.6 (h) cm) with divider panels that have open doors and a wire cup (9.5 (h) 7.6 (d) cm) in the two extreme chambers. In the second-stage (interaction stage), the experimental mouse was removed and a novel age- and sex- matched C57BL/6J mouse (social target) was placed under one wire cup and the experimental mouse was left to explore freely again. The area containing the novel mouse was defined as the social area, the area with the empty wire cup as the non-social area, and the middle as the neutral area. The following measures were recorded using Ethovision for statistical analyses: time spent in each

chamber, number of direct interactions between the experimental mouse and the wire cup (referred as nose-to-grid events; NTG), interaction time between the experimental mouse and the wire cup, total distance traveled, mean velocity and frequency of passes through the center chamber.

4. **Magnetic Resonance Imaging**

- **Acquisition**: Each mouse was weighed prior to each scan and anesthesia was induced with 3.5% isoflurane for at least 4 minutes. In vivo longitudinal T1-weighted structural MRI scans (FLASH; 100 $um^3$ voxels; matrix size 180 x 160 x 90, TR: 20ms, TE: 4.5 ms, flip angle: 10°), scan duration: 45 minutes) were acquired with a 7T Bruker Biospec 70/30 MRI scanner equipped with a cryogenically cooled surface coil at the Douglas Research Centre under a mixture of dexmedetomidine (0.25 mg/kg/h) and 1% isoflurane, in a 80% air and 20% oxygen mixture. Body temperature was maintained by hot air at 36.5℃. After each scan, mice recovered for at least 5 minutes on a heating pad under their home cage.
- **MRI Preprocessing:** Images were exported as Bruker format and converted to NIFTI using Brkraw (version 0.3.3), quality controlled (https://github.com/CoBrALab/documentation/wiki/Mouse-QC-Manual-(Structural)) and preprocessed. Preprocessing included denoising, N4 bias-field correction (Tustison et al., 2010) and affinely registered to an average mouse template using in-house pipeline developed at our laboratory (https://github.com/CoBrALab/documentation/wiki/Mouse-Scan-Preprocessing). Images were visually inspected for quality control prior to analyses.
- **Deformation-based morphometry:** Preprocessed images were used as an input for two-level deformation-based morphometry (https://github.com/CobraLab/twolevel_ants_dbm) (Germann et al., 2025). Using

registration tools from ANTs (https://github.com/ANTsX/ANTs) it evaluates longitudinal voxel-wise changes of brain volume using linear and nonlinear registration steps. In the first level, scans from all timepoints for a given subject were registered and used to create a subject-specific average, and in the second level the subject-specific averages were used to build a population-level average. In each level, images were aligned using affine transformations (3 translations, 3 rotations, scaling and shearing), followed by nonlinear transformations (Guma et al., 2023). Log-transformed Jacobian determinants are computed at the first level (Chung et al., 2001), resampled into the population-level average template and blurred at 0.2 mm full-width-at-half-maximum (FWHM). Voxels with positive values represent expansions relative to the population-level average, while negative values represent contractions. We used the Jacobian determinants only from the nonlinear transformations (referred to as relative Jacobians) for statistical analyses. By explicitly modeling only the non-linear part of the deformations, the relative Jacobian determinants remove residual global linear transformations (attributable to differences in total brain size, a major source of variability between mice (Lerch et al., 2012)) and allow us to examine relative volume changes to total brain volume.

5. **Partial Least Squares**

PLSC is a multivariate technique that discovers optimal weighted linear combinations between two sets of variables (McIntosh & Lobaugh, 2004; Zeighami et al., n.d.). Here, PLSC was performed on measures of brain volume and behavior (1) on within-subject change and (2) at post-CVS. For the first analysis on longitudinal change, we calculated the difference between post- and pre-CVS measures in brain volume (within-subject voxel-wise change;

https://github.com/CoBrALab/documentation/wiki/Create-a-nifti-of-within-subject-change-(2

-timepoints,-output-from-dbm)) and behavior. For the second analysis we used PLSC on cross-sectional measures of brain volume (voxel-wise relative Jacobian determinants) and behavior at the post-CVS timepoint.

First, a correlation matrix was computed from z-scored brain and behavior measures. Singular value decomposition (SVD) was applied to the brain-behavior matrix to yield a set of orthogonal latent variables (LVs). This generated singular values that describe the proportion of the variance explained by each LV, and a set of "brain weights" and "behavioral weights" describing how each voxel or behavioral variable weights onto a given LV. Brain and behavior weights were projected to the individual data to generate subject-specific brain and behavior scores. The statistical significance of each LV was assessed using permutation testing (n=1000). Here, the rows of the z-score brain matrix were randomly shuffled 1000 times to generate a null distribution of brain-behavior correlations. SVD was run on each of these correlations generating a null distribution of singular values, which was used to generate p-values. LVs were considered significant at $p<0.05$. The contribution of original brain and behavior variables to each LV was evaluated using bootstrap resampling (n=1000). Subjects were randomly sampled from both the brain and behavior matrices 1000 times. SVD was applied on each to generate a sampling distribution for each weight of the singular vectors. Bootstrap ratios were calculated as the ratio of each singular vector weight and its bootstrap-estimated standard error. These were thresholded at values corresponding to a 95% confidence interval. Voxels or behavioral variables that make higher contributions to the LV have high bootstrap ratios. Analyses were performed using Python version 3.6.8.

### 6. Structural connectivity

We used a graph theoretic framework to examine the topography of structural connections between regions that significantly covaried with anxiety- and depressive-like behavior (see

Partial Least Squares Correlation in the Multivariate analysis section). We leveraged the high-resolution models of the mouse connectome developed by Knox and colleagues (Knox et al., 2019). These models are based on the 428 viral injection experiments in C57BL/6J male mice obtained from the Allen Mouse Brain Connectivity Atlas (Oh et al., 2014). These experiments are a set of recombinant adeno-associated viruses that express green fluorescent protein and label monosynaptic connections. This data was registered onto the Allen Mouse Brain Common Coordinate Framework (Wang et al., 2020) and spatial connectivity at each voxel was inferred using a kernel-weighted average of the projection patterns of nearby injections. In this analysis, brain weights from PLS were thresholded at values corresponding to a 95% confidence interval and binarized. We then labeled groups of connected voxels (referred to as nodes throughout the remainder of this manuscript) and obtained a binarized structural connectivity matrix. We calculated the overlap between spatial connectivity patterns of each node as the proportion of voxels with tracer signal from node $x_1$ that overlap with the spatial distribution of node $x_2$. (For more details see Supplementary Figure 1). To evaluate the structural connectivity of voxels in the left hemisphere we flipped them along the sagittal midline. The final output was a directed and weighted adjacency matrix.

The significance of these overlaps was evaluated by building a null distribution of random connections to determine whether our groups of voxels were significantly connected. Here, we placed 1000 spheres at random locations in the brain (volume of the sphere was the average volume of the nodes for each analysis), obtained binarized maps of their structural connectivity and calculated their overlap to the empirical groups of voxels. (For more details see Supplementary Figure 1) as means of estimating p-values for each connection. After removing non-significant connections we binarized the adjacency matrix and computed the normalized rich club coefficient as in (Fulcher & Fornito, 2016). Specifically, this was defined as the ratio between the empirical rich-club coefficient and the mean rich-club coefficient of 1000 rewired networks. For each random network we maintained the same in- and out-degree distributions as implemented in the Brain Connectivity Toolbox (Rubinov & Sporns, 2010). We assessed statistical significance by obtaining p-values from this null distribution. We then used the Louvain algorithm to explore the presence of communities/modules in our directed connectome as in (Coletta et al., 2020). We varied the resolution parameter gamma (y) from 0.3 to 3.0 in 0.1 steps with 100 repetitions in each step, then performed consensus clustering and obtained community subdivisions of the connectome for each of the resolutions. We assessed the spatial similarity between the community partitions at each resolution using the adjusted mutual information and identified a gamma range with topographically stable community partitions as in (Coletta et al., 2020). To classify potential roles of our groups of voxels according to their intra- and inter-community connections we calculated the participation coefficient as in (Guimerà & Nunes Amaral, 2005) in which if a node has a participation coefficient closest to 1 its links are more distributed among all communities. Structural connectivity matrices were derived using Python version 3.8.2 and graph theory analyses were performed using the Brain Connectivity Toolbox (https://sites.google.com/site/bctnet/) on Matlab version 2020b.

### 7. Spatial gene expression

To investigate potential molecular mechanisms underlying the significant latent variables identified from our PLS analyses we used the Allen Institute's coronal mouse gene expression dataset in C57BL/6 male mice (post-natal day 56). In this dataset, brains were sliced in coronal sections of 200um width and gene expression was probed using in-situ hybridization. More details about this data set are found in (Lein et al., 2006). We used the coronal expression dataset containing a total of 4081 unique genes. For each gene, we downloaded the expression energy images at 200 um resolution. Each image was realigned to RAS+ orientation, origin translated inferior from bregma and converted to MINC format.

After removing non-brain related genes, we performed our analyses on 4071 normalized gene expression images (3836 unique genes). Normalization was done at each voxel by covarying for total expression across all genes. We first computed the Spearman correlation between spatial gene expression patterns and the brain weights from PLS, then ranked genes based on these correlations. We then identified gene ontology (GO) terms with genes that were preferentially distributed at the extreme ends of the ranked gene list. As in Ritchie et al (Ritchie et al., 2018), we removed very large and small modules, and retained those that contained between 10 and 200 genes (inclusive). We quantified enrichment as in (Hoops et al., 2024) in which, for a given module, we calculated the area under the curve (AUC) formed by the cumulative distribution of genes within a module as a function of rank. To test the statistical significance of AUC values, we computed 5000 random phenotypes, each generated from the average spatial gene expression of 10 random genes, and built a null distribution of AUC values. Finally, p-values obtained from testing multiple GO terms were FDR-corrected. Analyses were performed R version 3.5.1

### Supplementary Figures

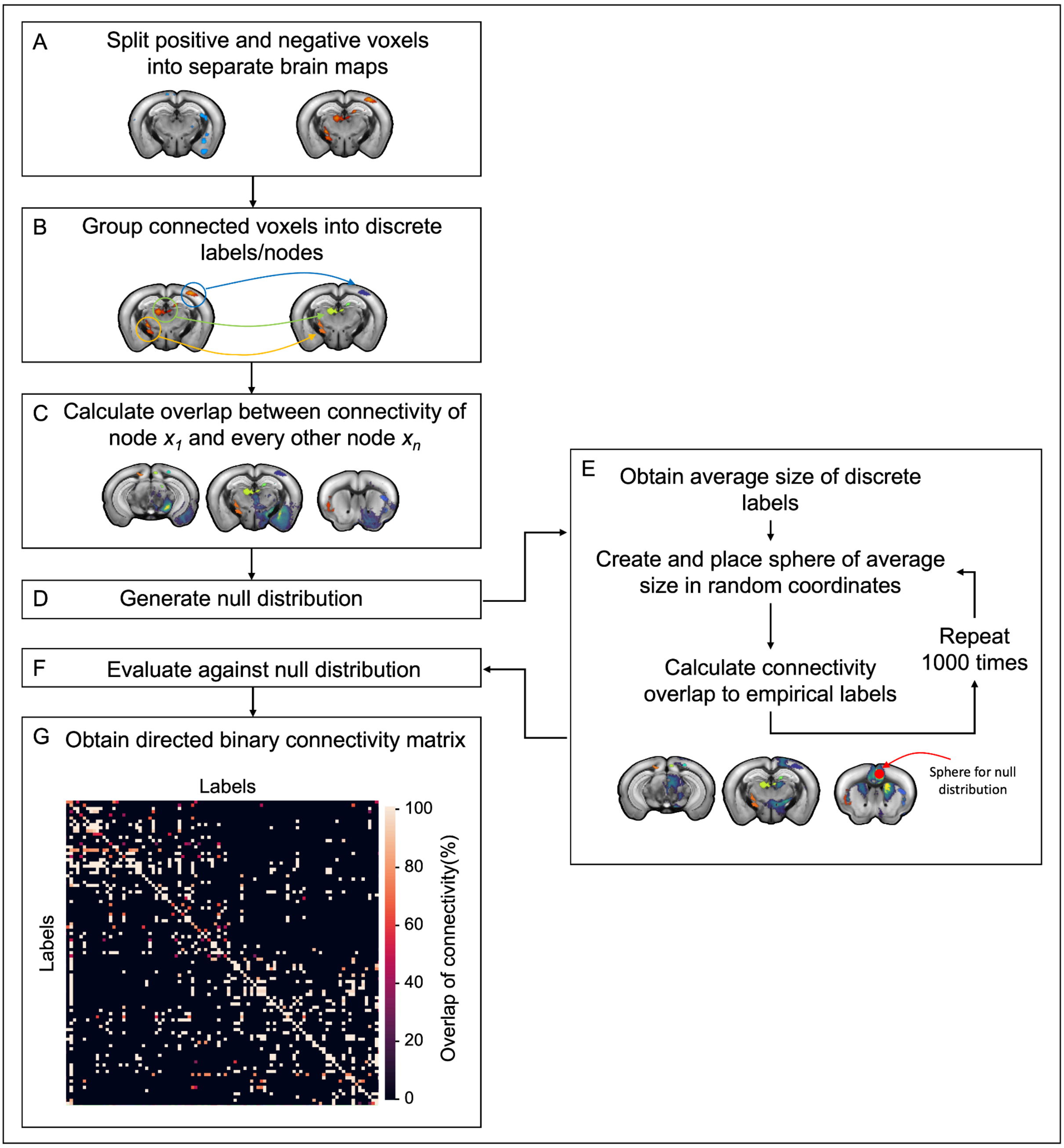


**Supplementary Figure 1. Steps used to create directed adjacency structural connectivity matrix of regions significantly associated with behavior, as derived from PLSC.** A) For our structural connectivity analysis we split the negative and the positive voxels into two separate brain maps, and performed our analyses separately for each. B) For each brain map, we grouped connected voxels into discrete labels (referred as nodes throughout the manuscript). C) For each node, we summed the structural connectivity spatial maps across all voxels in that node to derive the structural connectivity of that node to every other voxel in the brain. We then calculated the spatial overlap between each node's structural connectivity and the other voxels that encompassed the remaining nodes. We refer to this overlap as "Overlap of connectivity" which denotes that a node sends monosynaptic projections to another node. D) To determine the significance of our derived connectivity overlaps we generated a null distribution of random structural connections. E) We first obtained the average size across all nodes, created a sphere with a diameter of the average node size, placed that sphere in a random location and calculated its connectivity overlap with our empirical nodes. This process was repeated 1000 times to generate our null distribution. F and G) After evaluating against the null distribution, we obtained a directed adjacency matrix of structural connectivity between those nodes in the brain map.

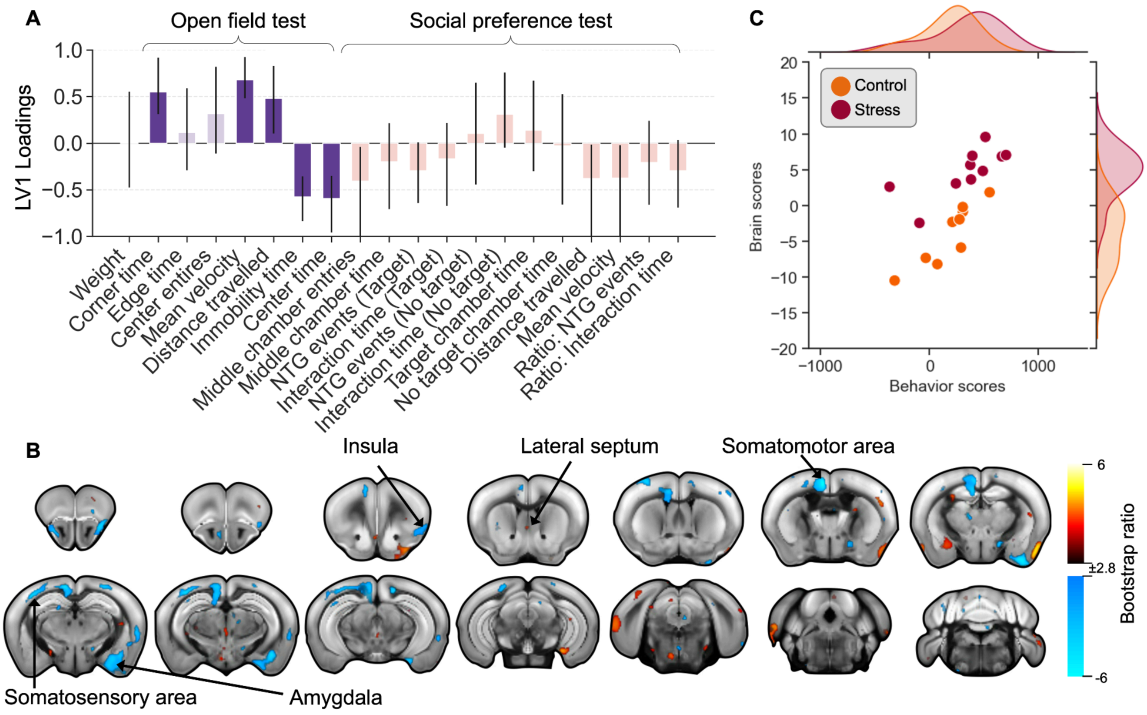


**Supplementary Figure 2. PLSC revealed signatures of stress-susceptibility at post-CVS time-point in females** (p=0.017, covariance explained: 20.76%). A) Weights of each behavioral variable onto its respective LV, indicating the correlation of each behaviour to the pattern. Size of the bar is estimated through singular value decomposition, confidence intervals are calculated by bootstrapping. Confidence intervals that cross the zero line should not be considered as a significant contribution to the LV. Anxiety-like behavior in purple and social withdrawal in orange. Changes in the open field and social preference test covary with the neuroanatomical changes denoted in (B). B) Weights of neuroanatomical changes onto its respective LV obtained from bootstrapping ratios overlaid on Allen CCFv3 template. Greater volume in warm colors (yellow-orange) and lower volume in cool colors (blue) indicate smaller volumes covary with behaviours. C) Subject-specific brain and behavior scores, color coded by group. (NTG: Nose-to-grid)

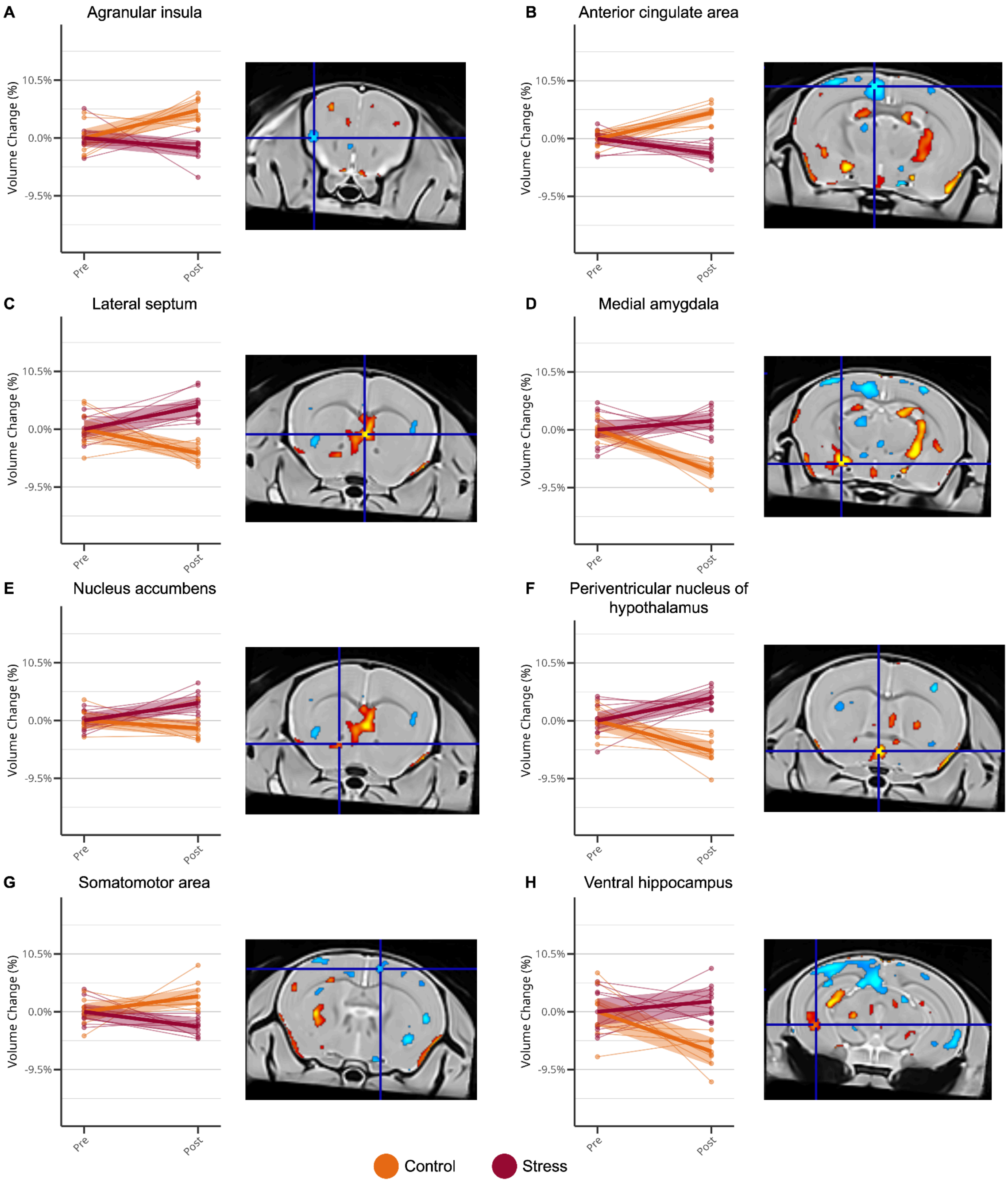


**Supplementary Figure 3. Peak voxels from linear mixed effects models examining group differences in volume change across time in female mice.** Plots of peak voxels (voxel showing largest effect within a cluster of voxels) from regions of interest illustrating percent volume change from baseline. Individual points represent observations and thin lines represent subject-level trajectories. Thick lines and shaded ribbons represent model-estimated means and 95% confidence intervals. Coronal brain slices with overlaid t-statistic maps (thresholded at 5% FDR) of group-by-timepoint interaction are shown to the left of each line plot. The blue cross-hair denotes the location of the peak voxel.

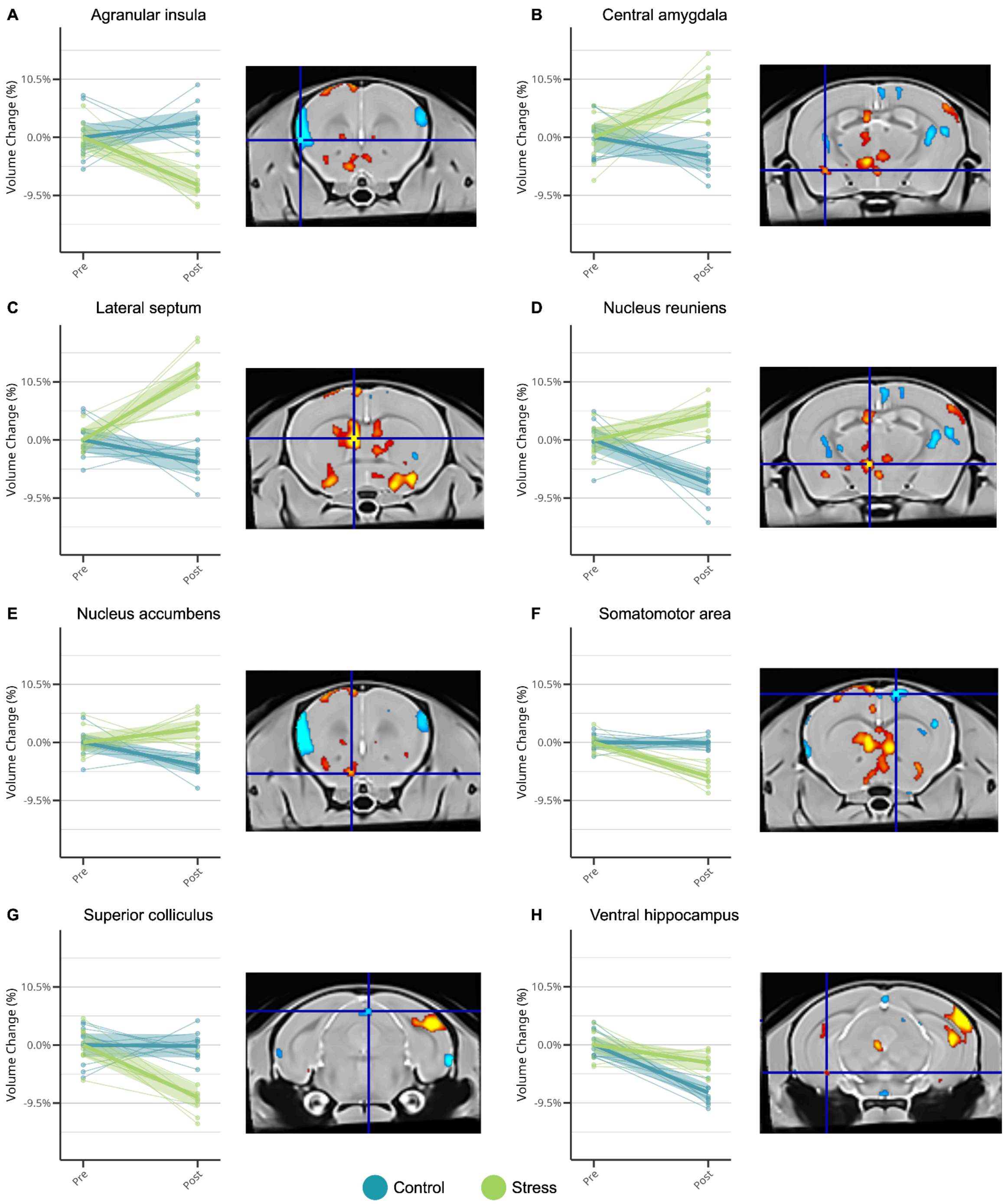


**Supplementary Figure 4. Line plots of peak voxels from linear mixed effects models examining group differences in volume change across time in male mice.** Plots of peak voxels (voxel showing largest effect within a cluster of voxels) from regions of interest illustrating percent volume change from baseline. Individual points represent observations and thin lines represent subject-level trajectories. Thick lines and shaded ribbons represent model-estimated means and 95% confidence intervals. Coronal brain slices with overlaid t-statistic maps (thresholded at 5% FDR) of group-by-timepoint interaction are shown to the left of each line plot. The blue cross-hair denotes the location of the peak voxel.

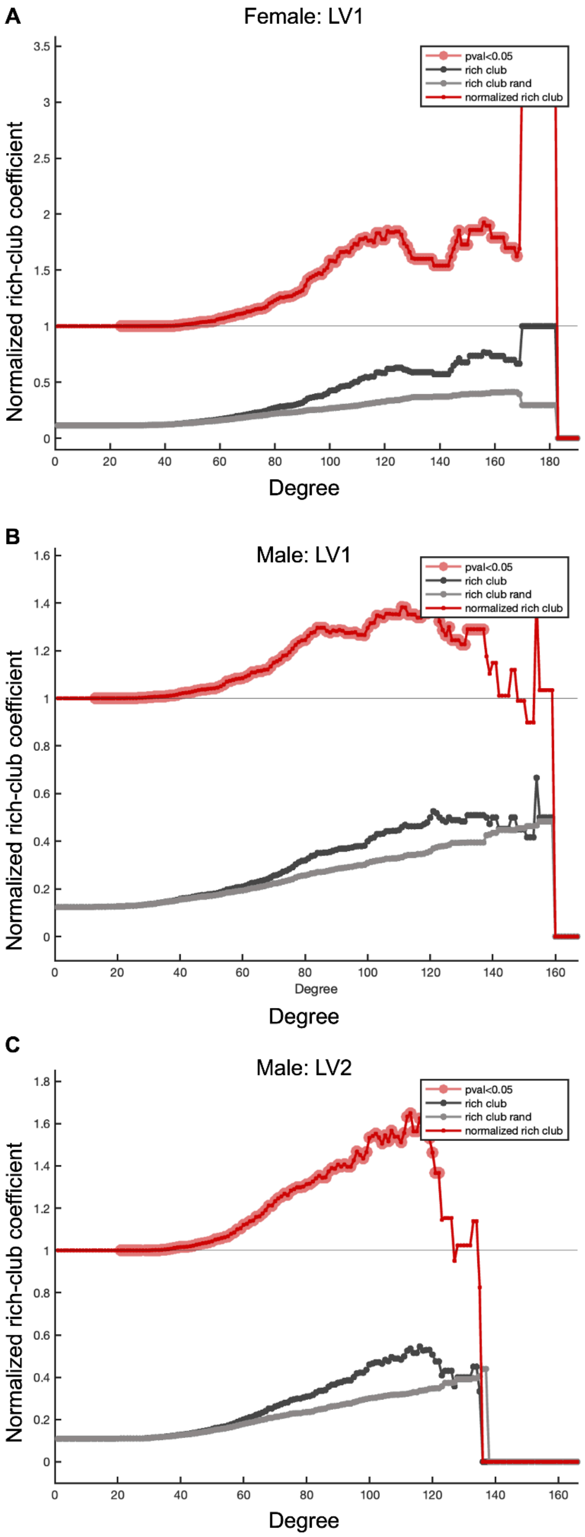


**Supplementary Figure 5. Topological rich club regime of structural connectomes underlying stress susceptibility signatures in female and male mice.** Normalized rich club coefficient (red) as a function of degree for significant latent variables from PLSC analysis in females (A) and males (B and C). Red circles indicate values that significantly higher that a null distribution of 1000 rewired networks.

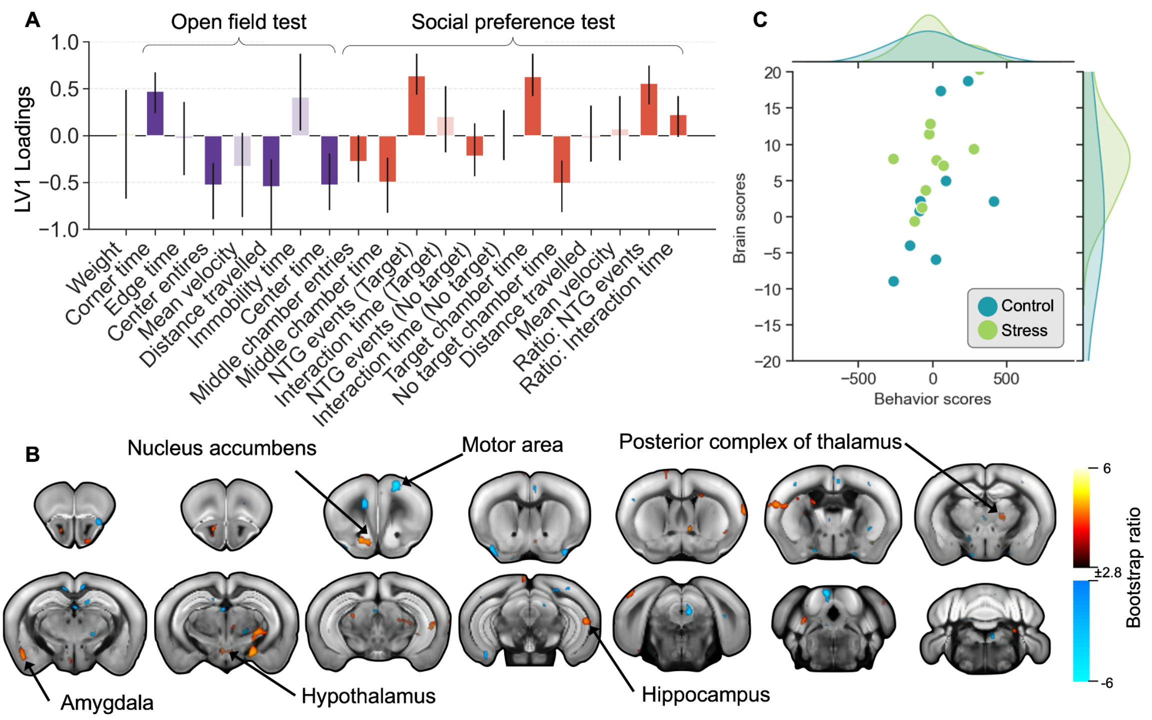


**Supplementary Figure 6. Non-significant covarying brain-behavior pattern of stress susceptibility on the change across time in male mice.** (p=0.28, covariance explained: 23.8%). A) Weights of each behavioral variable onto its respective LV, indicating the correlation of each behaviour to the pattern. Size of the bar is estimated through singular value decomposition, confidence intervals are calculated by bootstrapping. Confidence intervals that cross the zero line should not be considered as a significant contribution to the LV. Anxiety-like behavior in purple and social withdrawal in orange. Changes in the open field and social preference test covary with the neuroanatomical changes denoted in (B). B) Weights of neuroanatomical changes onto its respective LV obtained from bootstrapping ratios overlaid on Allen CCFv3 template. Greater volume in warm colors (yellow-orange) and lower volume in cool colors (blue) indicate smaller volumes covary with behaviours. C) Subject-specific brain and behavior scores, color coded by group. (NTG: Nose-to-grid)

### Supplementary Tables

**Supplementary Table 1.** Acronyms and label names of annotations used in analyses.

| Acronym | Label name |
|---|---|
| ACA | Anterior cingulate area |
| AI | Agranular insular area |
| AN | Ansiform lobule |
| AT | Anterior tegmental nucleus |
| AUD | Auditory areas |
| BLA | Basolateral amygdalar nucleus |
| BMA | Basomedial amygdalar nucleus |
| CENT | Central lobule |
| CLA | Claustrum |
| COPY | Copula pyramidis |
| CTXpl | Cortical plate |
| CTXsp | Cortical subplate |
| CUL | Culmen |
| CUN | Cuneiform nucleus |
| DEC | Declive (VI) |
| DN | Dentate nucleus |
| DORpm | Thalamus, polymodal association cortex related |
| DORsm | Thalamus, sensory-motor cortex related |
| ECT | Ectorhinal area |
| EP | Endopiriform nucleus |
| EW | Edinger-Westphal nucleus |
| FL | Flocculus |
| FN | Fastigial nucleus |
| FOTU | Folium-tuber vermis (VII) |
| FRP | Frontal pole, cerebral cortex |
| GU | Gustatory areas |
| HEM | Hemispheric regions |
| HPF | Hippocampal formation |
| HY | Hypothalamus |
| IC | Inferior colliculus |
| III | Oculomotor nucleus |
| ILA | Infralimbic area |
| IP | Interposed nucleus |
| Isocortex | Isocortex |
| IV | Trochlear nucleus |

| | |
|---|---|
| LA | Lateral amygdalar nucleus |
| LING | Lingula (I) |
| LSX | Lateral septal complex |
| LT | Lateral terminal nucleus of the accessory optic tract |
| LZ | Hypothalamic lateral zone |
| MBmot | Midbrain, motor related |
| MBsen | Midbrain, sensory related |
| MBsta | Midbrain, behavioral state related |
| ME | Median eminence |
| MEV | Midbrain trigeminal nucleus |
| MEZ | Hypothalamic medial zone |
| MO | Somatomotor areas |
| MRN | Midbrain reticular nucleus |
| MY | Medulla |
| MY-mot | Medulla, motor related |
| MY-sat | Medulla, behavioral state related |
| MY-sen | Medulla, sensory related |
| NB | Nucleus of the brachium of the inferior colliculus |
| NOD | Nodulus (X) |
| OLF | Olfactory areas |
| ORB | Orbital area |
| P | Pons |
| P-mot | Pons, motor related |
| P-sat | Pons, behavioral state related |
| P-sen | Pons, sensory related |
| PA | Posterior amygdalar nucleus |
| PAG | Periaqueductal gray |
| PAL | Pallidum |
| PALc | Pallidum, caudal region |
| PALd | Pallidum, dorsal region |
| PALm | Pallidum, medial region |
| PALv | Pallidum, ventral region |
| PBG | Parabigeminal nucleus |
| PERI | Perirhinal area |
| PFL | Paraflocculus |
| PL | Prelimbic area |
| PPN | Pedunculopontine nucleus |
| PRM | Paramedian lobule |
| PRT | Pretectal region |
| PTLp | Posterior parietal association areas |
| PVR | Periventricular region |

| | |
|---|---|
| PVZ | Periventricular zone |
| PYR | Pyramus (VIII) |
| RAmb | Midbrain raphe nuclei |
| RN | Red nucleus |
| RR | Midbrain reticular nucleus, retrorubral area |
| RSP | Retrosplenial area |
| SAG | Nucleus sagulum |
| sAMY | Striatum-like amygdalar nuclei |
| SCm | Superior colliculus, motor related |
| SCs | Superior colliculus, sensory related |
| SIM | Simple lobule |
| SNc | Substantia nigra, compact part |
| SNr | Substantia nigra, reticular part |
| SS | Somatosensory areas |
| STR | Striatum |
| STRd | Striatum dorsal region |
| STRv | Striatum ventral region |
| TEa | Temporal association areas |
| TH | Thalamus |
| UVU | Uvula (IX) |
| VERM | Vermal regions |
| VIS | Visual areas |
| VISC | Visceral area |
| VTA | Ventral tegmental area |
| VTN | Ventral tegmental nucleus |
